# PureseqTM: efficient and accurate prediction of transmembrane topology from amino acid sequence only

**DOI:** 10.1101/627307

**Authors:** Qing Wang, Chong-ming Ni, Zhen Li, Xiu-feng Li, Ren-min Han, Feng Zhao, Jinbo Xu, Xin Gao, Sheng Wang

## Abstract

**Motivation:** Rapid and accurate identification of transmembrane (TM) topology is well suited for the annotation of the entire membrane proteome. It is the initial step of predicting the structure and function of membrane proteins. However, existing methods that utilize only amino acid sequence information suffer from low prediction accuracy, whereas methods that exploit sequence profile or consensus need too much computational time.

**Method:** Here we propose a deep learning framework DeepCNF that predicts TM topology from amino acid sequence only. Compared to previous sequence-based approaches that use hidden Markov models or dynamic Bayesian networks, DeepCNF is able to incorporate much more contextual information by a hierarchical deep neural network, while simultaneously modeling the interdependency between adjacent topology labels.

**Result:** Experimental results show that PureseqTM not only outperforms existing sequence-based methods, but also reaches or even surpasses the profile/consensus methods. On the 39 newly released membrane proteins, our approach successfully identifies the correct TM segments and boundaries for at least 3 cases while all existing methods fail to do so. When applied to the entire human proteome, our method can identify the incorrect annotations of TM regions by UniProt and discover the membrane-related proteins that are not manually curated as membrane proteins.

**Availability:** http://pureseqtm.predmp.com/

## Introduction

Transmembrane proteins (TMPs) are key players in energy production, material transport, and communication between cells [1]. TMPs are encoded by ∼30% genes in the various genomes [2] and have been targeted by ∼50% of therapeutic drugs [3]. Despite their abundance and importance, the number of solved TMPs structures is relatively low compared to that of non-transmembrane proteins (non-TMPs). In particular, under the 40% sequence identity cutoff, there are only about 1500 non-redundant TMPs whereas the number of non-redundant non-TMPs is more than 34000. The underlying reason is that the experimental determination of TMPs is challenging as membrane proteins are often too large for NMR spectroscopy and difficult to be crystallized for X-ray crystallography [4]. Thus, it is critical to develop computational methods for the prediction of TMP structures from amino acid sequences, and the initial step is the accurate identification of the transmembrane topology [5].

As shown in the left part of Figure 1, transmembrane (TM) topology refers to the locations of the membrane-spanning segments, which could be represented as a 1D 0/1 string to indicate the location of each residue to reside in (label 1) or out of (label 0) the membrane. This simple but direct definition of TM topology is consistent with the 3-label definition used by many other works that divide non-TM regions (i.e., label 0) into inner or outer classes [6–10]. In this work, we only focus on the prediction of TM topology of the alpha-helical TMPs because of the following two facts: (i) almost all the TM regions in Eukaryotic TMPs are alpha-helical except some beta-barrel in the mitochondrial membrane [8]; (ii) more than 85% of the available TMPs that have 3D structures belong to the alpha-helical class [11]. If there is only one TM segment, then this membrane protein is denoted as a single-pass transmembrane protein (sTMP); similarly, a multi-pass transmembrane protein (mTMP) will contain two or more TM segments. Here we mainly focus on the topology prediction of multi-pass transmembrane proteins as about 75% of the current available alpha-helical TMPs are mTMPs [11].

**Figure 1.**
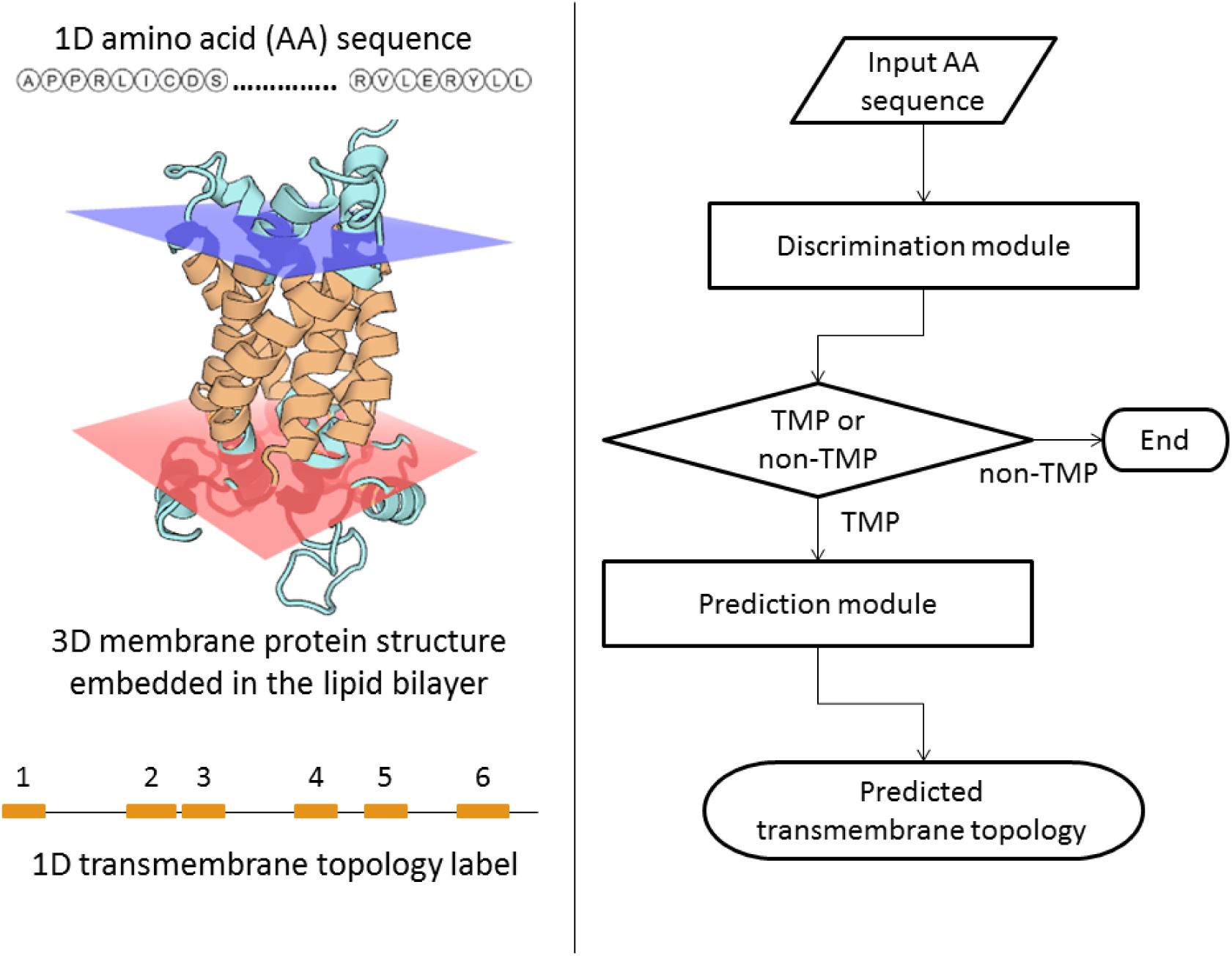
Illustration of the transmembrane (TM) topology of a multi-pass transmembrane protein and our proposed model for the prediction of TM topology. **Left**: the 1D amino acid sequence of an alpha-helical membrane protein (PDB ID: 2lckA) folds into a 3D structure embedded in the lipid bilayer membrane, in which those residues embedded are denoted as the TM topology. **Right**: our proposed model that consists of two modules, where the first module is for discriminating transmembrane proteins (TMPs) and non-TMPs, and the second module is for predicting the TM topology.

Till now, a variety of approaches have been proposed to predict the 1D TM topology from the input sequence of a membrane protein. These approaches can be roughly categorized into three classes: (a) single-sequence-based methods that only rely on the input amino acid sequence information (or, ‘pureseq’ features). Two representative methods are TMHMM/Phobius [7, 12] and Philius [8], where the former established a hidden Markov model (HMM) and the latter employed a dynamic Bayesian network (DBN) model. The advantage of the pureseq methods is their fast running speed, while the disadvantage is the relatively low prediction accuracy; (b) evolutionary-based methods that consider the information embedded in the homologous multiple sequence alignment (MSA) through evolutionary analysis (or, ‘profile’ features). Two representative methods are OCTOPUS [10] and MEMSAT-SVM [9]. The advantage of the profile methods is their improved prediction accuracy over pureseq methods, but at the cost of significantly reduced running speed due to the search for MSA; (c) consensus methods that combine the outputs from different predictors. Topcons2 [13] and CCTOP [6] are the two representatives. As the consensus methods also integrate the profile methods, their running speed cannot compete with that of the pureseq methods. Thus, it remains a question of if we can develop an approach for TM topology prediction that can reach the accuracy of profile or consensus methods but as efficient as pureseq methods.

Recently, we developed a deep learning framework deep convolutional neural fields (DeepCNF) [14] for a variety of protein sequence labeling problems ranging from secondary structure element (SSE) [15], solvent accessibility (ACC) [16], to order/disorder region (DISO) [17, 18], which obtained the state-of-the-art performance according to the third-party evaluations [19–22]. In this work, we employed DeepCNF for the prediction of TM topology labels based on amino acid sequence information only (denoted as PureseqTM). Briefly, DeepCNF can be viewed as conditional random field (CRF) with deep convolutional neural network (DCNN) as its non-linear feature generating function. DeepCNF can model not only complex relationship between the input features and TM labels, but also the correlation among adjacent TM labels. These properties give DeepCNF better ability to model long-range dependencies embedded in the input features, and better performance over CRF (a model compatible to or even better than HMM and DBN) and DCNN on 1D sequence labeling tasks. Furthermore, besides considering amino acids as simply 20 alphabets, we may also take into account the physical-chemical properties of amino acids [23]. Consequently, our proposed method, PureseqTM, can easily take as input these pureseq features, and achieve the performance compatible to or even better than profile and consensus methods, while keeping the similar running speed of pureseq methods.

As shown in the right part of Figure 1, PureseqTM has two modules: (i) a module for discriminating TMPs and non-TMPs, and (ii) a module for predicting TM topology. Experimental results show that PureseqTM greatly outperforms existing pureseq methods Phobius and Philius, especially on the identification of the correct number and boundaries of the TM segments (measured by protein-level and segment-level accuracy, respectively). Specifically, on the 39 newly released mTMPs, PureseqTM achieved the best performance 0.667 in terms of protein-level accuracy, which is 12.9%, 7.7% and 5.2% better than Phobius, Philius and even Topcons2, respectively. Moreover, PureseqTM correctly identified the number and boundaries of the TM segments for at least 3 cases among this dataset where Phobius, Philius, or Topcons2 was not able to do so. Finally, we applied our method on the entire human proteome from UniProt [24]. The results indicate that PureseqTM can not only identify the incorrect annotations of TM regions by UniProt, but also discover the membrane-related proteins that are not reviewed as membrane proteins by UniProt. Since it is time-consuming to generate sequence profiles, our proposed method is a useful tool for proteome-wide TM topology prediction.

## Results

### Model Architecture

In general, as shown in Figure 2, the model architecture of PureseqTM could be considered as an integration of HMM and DBN to produce an initial estimation of the probabilities of the transmembrane topology labels as well as the probabilities of their transitions (or more precisely, topology change). Then the long-range dependencies embedded in the information are in turn effectively exploited by a DeepCNF model to predict the binary topology labels at each amino acid position.

**Figure 2.**
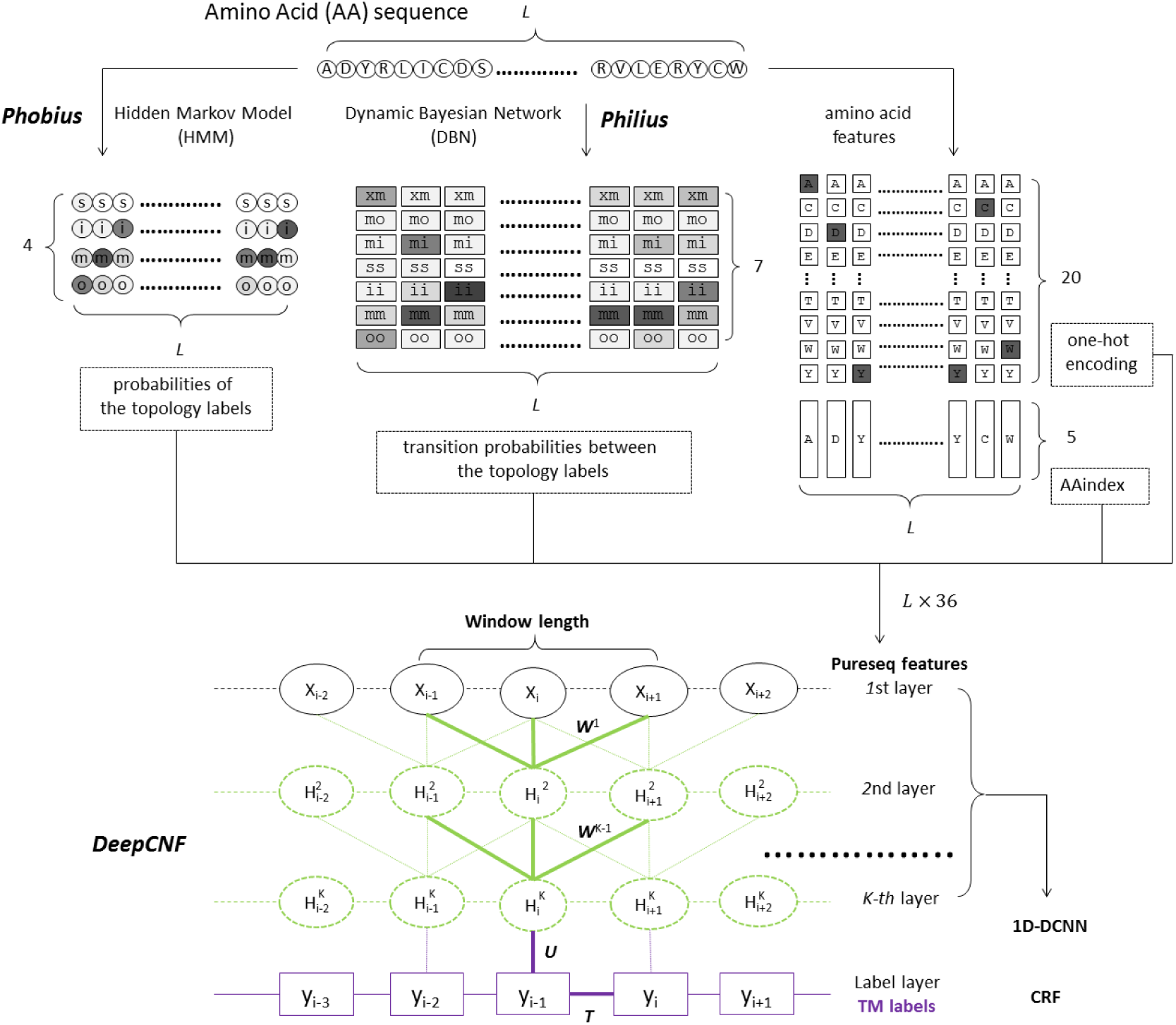
Overview of our Deep Convolutional Neural Fields (DeepCNF) model for transmembrane (TM) topology label prediction from the amino acid sequence features only (i.e., pureseq features). Here *L* is the amino acid sequence length of the input protein. The probabilities of the output of Hidden Markov Model (HMM) and Dynamic Bayesian Network (DBN) are displayed in gray scale, where darker or lighter indicates higher or lower probabilities, respectively. CRF denotes Conditional Random Field (in purple), and 1D-DCNN denotes 1D Deep Convolutional Neural Network (in light green).

#### HMM features generated by Phobius

Phobius [12] considers the transmembrane topology prediction problem as a supervised learning problem over the observed input amino acid sequences ***a***= *a*_1_,…,*a_L_*, and output of the hidden topology labels ***y*** = *y*_1_,…,*y_L_*. The *y_i_*is from the four topology labels {`s,i,m,o`}, which correspond respectively to signal peptides, cytoplasmic (‘inside’) loops, membrane-spanning segments, and non-cytoplasmic (‘outside’) loops. Then Phobius established an HMM that is a generative model to produce a joint probability distribution between the observed sequence ***a*** and the hidden states ***y***. When the HMM model is trained, it is possible to infer the posterior probability of the four topology labels at each amino acid position via a forward-backward algorithm. Thus, the generated four probabilities of the topology labels from Phobius become the HMM features for our proposed DeepCNF model.

#### DBN features generated by Philius

DBN could be regarded as a strict generalizations of HMM [25], where the latter only contains two variables (one observation and one hidden state) in each time frame i and one connection between adjacent frames (i.e., the connection between the adjacent hidden states). On the contrary, DBN can model multiple variables in each frame as well as more than one connection between adjacent frames. This property enables DBN to encode the states that depict the transitions between the adjacent topology labels (i.e., ‘`changeState`’ according to Philius [8]). In particular, `changeState` can take value 0 or 1. If 1 is taken, then the topology label in the next frame will be changed. Therefore, instead of predicting the posterior probability of the four topology labels (i.e., `changeState` is set to 0) which respectively correspond to ‘`ss`’, ‘`ii`’, ‘`mm`’, and ‘`oo`’, Philius is able to predict additional four posterior probability of the topology label transitions (i.e., `changeState` is set to 1) which respectively correspond to ‘`xs`’, ‘`xi`’, ‘`xm`’ and ‘`xo`’ where ‘`x`’ indicate any other topology label. However, as the signal peptide starts from the N-terminal and the probability of ‘`xs’` only has a value at the first frame, we may delete this state. Also, the ‘`x`’ could only be ‘`m`’ for ‘`xi`’ and ‘`xo`’ because of the state transition diagram in topology prediction. In summary, the generated seven transition probabilities between the topology labels from Philius become the DBN features for our proposed DeepCNF model.

#### Additional amino acid features

In addition to the predicted probabilities of the topology labels from Phobius and Philius, we further utilize the features embedded in the input amino acid sequence (i.e., ‘pureseq’ features). One straightforward approach (denoted as one-hot encoding) uses a binary vector of 20 elements to indicate the amino acid type at position i. However, the 20 amino acids are not simply alphabetic letters, as they encode a variety of physiochemical properties. These properties of the 20 amino acids could be obtained from an on-line database (AAindex) [23] that forms a 20X494 matrix (i.e., each amino acid has 494 physiochemical properties). This high dimensional and redundant data can be reduced to a 20×5 matrix (i.e., each amino acid can be represented by a 5-dimentional vector), which represents bipolar, secondary structure, molecular volume, relative amino acid composition, and electrostatic charge, respectively [26]. Consequently, these 20+5 pureseq features are added to our proposed DeepCNF model.

#### Binary topology label prediction by DeepCNF

It has been reported that the DeepCNF model was successfully applied to a variety of sequence labeling problems, such as protein secondary structure prediction [15], protein order/disorder region prediction [18], and detecting the boundaries of expressed transcripts from RNA-seq reads alignment [27]. Generally speaking, DeepCNF has two modules, CRF and DCNN. DeepCNF can not only model complex relationship between the features from the amino acid sequence and the topology label by a deep hierarchical architecture that allows the model to capture long-range dependencies embedded in these features (in parameters ***W*** and ***U***), but also explicitly depict the interdependency between adjacent topology labels (in parameter ***T***).

### Performance evaluation

We measure the prediction results in terms of the following evaluation criteria: protein-level accuracy, segment-level accuracy, and residue-level accuracy. The protein-level accuracy (pAccu) refers to the definition that a correct prediction of the whole protein should have the correct number of transmembrane segments at approximately correct locations (overlap of at least five residues) [13]. For segment-level accuracy, we use segment recall (sReca) and segment precision (sPrec). The segment recall is defined as the approximately correct prediction of the transmembrane segment with overlap of at least five residues to the ground-truth segment; similarly, the definition for segment precision is the approximately correct ground-truth of the transmembrane segment with overlap of at least five residues to the predicted segment [8].

The residue-level accuracy is defined at each residue, which consists of Q2 accuracy, SOV score, recall (Reca), precision (Prec) and Matthews correlation coefficient (Mcc), respectively. The Q2 accuracy is defined as the percentage of residues for which the predicted transmembrane topology label is correct. The Segment OVerlap (SOV) score [28] measures how well the ground-truth and the predicted transmembrane regions match, especially at the middle region instead of terminal regions (see Section S1 in Supplemental Material for details). In order to calculate recall, precision and Mcc, we define true positives (TP) and true negatives (TN) as the numbers of correctly predicted transmembrane and non-transmembrane residues, respectively; whereas false positives (FP) and false negatives (FN) are the numbers of misclassified transmembrane and non-transmembrane residues, respectively. Then recall and precision is defined as TP = (TP + FN) and TP = (TP + FP), respectively. Mcc is defined as follows:

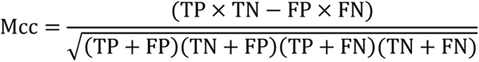

### Performance on validation dataset

The performance of our approach PureseqTM was compared to the performance of Phobius [12] and Philius [8] on the 164 validation dataset (see Method for details on how we create the 164 training and 164 validation dataset). Furthermore, to discover the importance of each input features, we conduct a study by incrementally adding the features from one-hot encoding, AAindex, HMM to DBN, respectively. This feature incremental study is critical as the HMM and DBN features originate from Phobius and Philius, and we need to show the performance to what extend PureseqTM can reach without using these features.

As shown in Table 1, all our feature combination strategies outperform Phobius and Philius in terms of the segment-level, and the majority of the residue-level accuracy, such as Q2, SOV, and Mcc. Notably, our DeepCNF model trained by only one-hot encoding features outperforms Phobius and Philius, especially in terms of Q2, SOV, and Mcc from the residue-level accuracy, and segment recall and precision. These results suggest that there exist some long-range dependencies between amino acid sequence and transmembrane topology, and this information can be learned by our deep learning model better than that by HMM and DBN. Furthermore, with more features being added to DeepCNF, the performance, especially in terms of protein-level accuracy, increases incrementally. When all features are added, our method can reach 0.573 protein-level accuracy, which is compatible to Philius.

**Table 1.** Overall transmembrane topology prediction accuracy on 164 membrane proteins from the validation dataset.

|  | pAccu | sReca | sPrec | Q2 | SOV | Reca | Prec | Mcc |
| --- | --- | --- | --- | --- | --- | --- | --- | --- |
| Phobius | 0.500 | 0.889 | 0.905 | 0.816 | 0.837 | 0.797 | 0.715 | 0.601 |
| Philius | 0.579 | 0.917 | 0.914 | 0.828 | 0.859 | 0.833 | 0.724 | 0.627 |
| DeepCNF trained by amino acid sequence only |  |  |  |  |  |  |  |  |
| One-hot | 0.543 | 0.935 | 0.917 | 0.836 | 0.865 | 0.786 | 0.785 | 0.648 |
| + AAindex | 0.561 | 0.945 | **0.922** | 0.838 | 0.868 | 0.796 | **0.789** | 0.653 |
| + HMM | 0.567 | **0.950** | 0.921 | 0.839 | 0.870 | **0.819** | 0.766 | 0.654 |
| + DBN | **0.579** | 0.935 | 0.912 | **0.842** | **0.881** | 0.804 | 0.771 | **0.654** |
| DeepCNF trained by sequence profile |  |  |  |  |  |  |  |  |
| Profile | 0.708 | 0.927 | 0.932 | 0.867 | 0.906 | 0.811 | 0.808 | 0.699 |
| + One-hot | 0.701 | 0.942 | 0.930 | 0.866 | 0.906 | 0.815 | **0.816** | 0.702 |
| + AAindex | **0.750** | **0.950** | **0.940** | **0.869** | **0.912** | 0.829 | 0.811 | **0.709** |
| + HMM | 0.732 | 0.952 | 0.938 | 0.866 | 0.911 | **0.830** | 0.808 | 0.705 |
| + DBN | 0.695 | 0.944 | 0.933 | 0.865 | 0.906 | 0.824 | 0.806 | 0.701 |

### Performance on test dataset

To show the performance on “real-world” cases, we challenged our method on the 39 mTMPs dataset, in which all data are released after Jul 2016 [11] and no homologous to the entries in the 328 training and validation set. Moreover, we performed a 3D structure comparison [29] between the proteins in the test dataset and those in the training dataset. Among the 39 mTMPs, 7 (15) of them have structural analogs in the 328 dataset whose TMscore > 0.65 (0.55). This means that at least 60% to 82% of the mTMPs in our test dataset are novel fold to the training dataset [30]. This resembles actual challenges that no sequence or structure similarity could be found for those newly obtained mTMPs.

For this task, we not only compared our PureseqTM with Phobius and Philius that rely on the input amino acid sequence only (i.e., ‘pureseq’ methods), but also compared with Topcons2 [13] which is a consensus approach relying on the evolutionary profile additionally. As shown in Table 2, not surprisingly, our method again achieves better performance than Phobius and Philius in terms of the key measurements in residue-level accuracy, such as Q2 (improved by 2.2%), SOV (improved by 3.6%) and Mcc (improved by 3.2%). Figure 3 shows the head-to-head comparison of the Mcc and SOV values between PureseqTM and the other three methods. These results indicate that PureseqTM outperforms others in terms of Mcc and SOV for a large portion of the cases in the test dataset.

**Figure 3.**
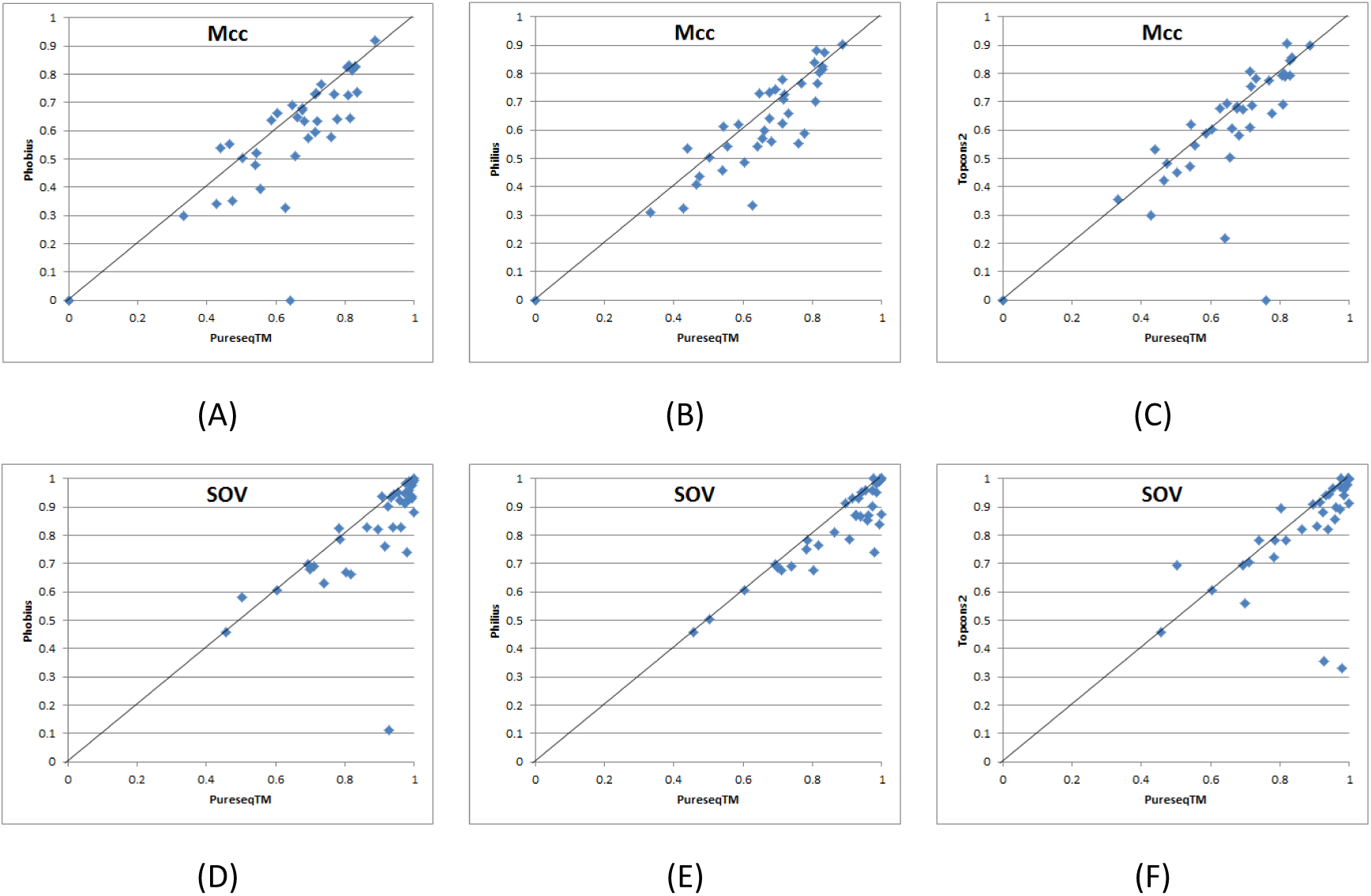
Quality comparison of the prediction by our method PureseqTM, with Phobius, Philius and Topcons2 on the 39 test dataset. (A) to (C): comparison between our method (X-axis) and other three methods (Y-axis) in terms of Matthews correlation coefficient (Mcc). (D) to (F): comparison between our method (X-axis) and other three methods (Y-axis) in terms of segment overlap score (SOV).

**Table 2.** Overall transmembrane topology prediction accuracy on 39 membrane proteins from the test dataset.

|  | pAccu | sReca | sPrec | Q2 | SOV | Reca | Prec | Mcc |
| --- | --- | --- | --- | --- | --- | --- | --- | --- |
| Phobius | 0.538 | 0.833 | 0.911 | 0.808 | 0.826 | 0.750 | 0.715 | 0.579 |
| Philius | 0.590 | 0.863 | **0.939** | 0.816 | 0.849 | **0.782** | 0.737 | 0.603 |
| Topcons2 | 0.615 | 0.851 | 0.910 | 0.819 | 0.840 | 0.759 | 0.724 | 0.593 |
| PureseqTM | **0.667** | **0.892** | 0.934 | **0.838** | **0.885** | 0.762 | **0.777** | **0.635** |
| PureseqTM<sup>P</sup> | 0.667 | 0.903 | 0.933 | 0.851 | 0.902 | 0.782 | 0.790 | 0.661 |

'P' indicates that the sequence profile feature is used.

It is surprising that PureseqTM achieved the best performance 0.667 in terms of protein-level accuracy, which is 12.9%, 7.7% and 5.2% better than Phobius, Philius and even Topcons2, respectively. These results show that the improvement of our method is even higher than that in the 164 validation set, especially in the protein-level accuracy. One possible explanation is that some of those mTMPs in the validation set overlap with the training data of Phobius and Philius. Consequently, these results indicate that our approach could be applied to a newly released mTMP that possibly has novel transmembrane topology, with only amino acid sequence as input.

Table 2 also shows a strange phenomenon that in terms of segment-level and residue-level accuracy, Philius, a pureseq method, outperforms the consensus method Topcons2 that relies on the evolutionary profile as well as the output of Philius. In order to explain this case, we conduct a similar feature incremental study on training data but start from the 40 evolution-related features (see Method for details of how to generate these profile features). As shown in the bottom part of Table 1, the performance reaches its peak till the AAindex features are added, and it decreases when adding HMM or DBN. This experiment might explain why the performance of Topcons2 is not comparable to that of Philius.

If the DeepCNF model trained by the sequence profile (i.e., profile model) is applied to the 39 test set, we find that in terms of residue-level accuracy, such as Q2 and SOV, the profile model gains ∼1-2% compared to the model trained by residue-related features (i.e., pureseq model). However, it seems that the protein-level and segment-level accuracy of the profile model does not improve much over that of the pureseq model, which might indicate that the evolution-related features could improve the boundary detection, but are not that useful for identifying the transmembrane segment. This hypothesis could also be used to explain the phenomenon why Topcons2 does not outperform Philius in terms of the protein-level and segment-level accuracy.

### Case studies of the test set

To further demonstrate the performance of PureseqTM, we select three case studies from the test set where the ground-truth of the transmembrane topology is known from PDBTM [11].

#### 5lnko

This protein is the chain o from the ovine respiratory complex I, which has 120 residues and 2 transmembrane helices. Note that this protein is structurally dissimilar to the training dataset, where the most similar protein only has TM-score 0.47 to this protein. This indicates that 5lnko has a novel fold. As shown in Figure 4, our method predicted the correct number of transmembrane helices, while Phobius and Philius predicted one, and Topcons2 failed to predict this protein as a TMP. In terms of the residue-level accuracy (Table 3), our method reached 0.806, 0.833 precision, 0.758 Mcc, 0.908 Q2, and 0.978 SOV, which indicated that the overlap between our prediction and the ground-truth is very large. Therefore, the success of this case indicates that PureseqTM could be applied to predict the transmembrane topology for those “real-world” new cases that no sequence or structure similarity could be found in the sequence or template database.

**Figure 4.**
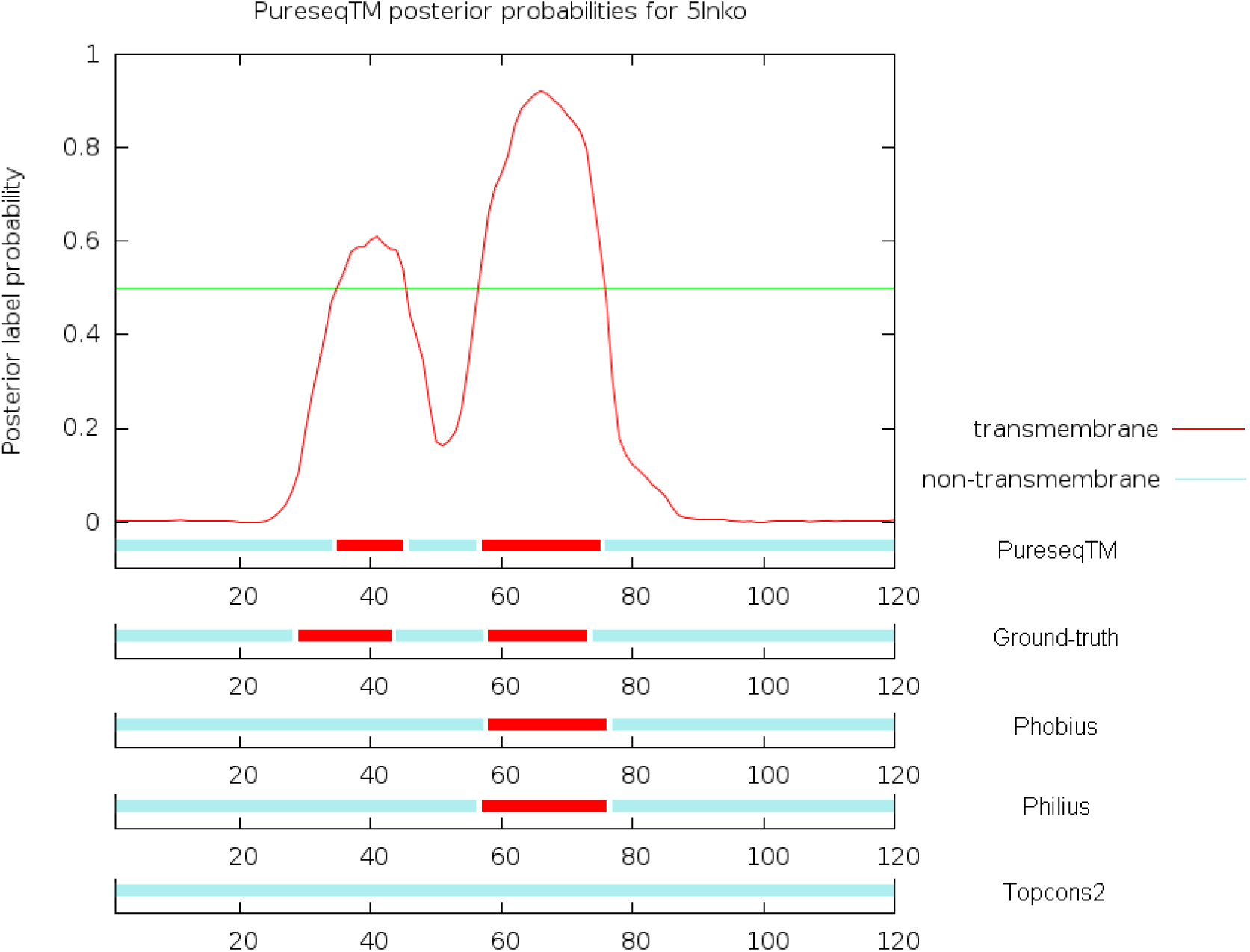
Case study of the transmembrane topology prediction of 5lnko. Here transmembrane or non-transmembrane regions are shown in red or cyan, respectively. The posterior probabilities generated by PureseqTM are shown in red curve, and the 0.5 threshold is shown in green line. (the same as below)

**Table 3.** Transmembrane topology prediction accuracy of 5lnko.

|  | pAccu | sReca | sPrec | Q2 | SOV | Reca | Prec | Mcc |
| --- | --- | --- | --- | --- | --- | --- | --- | --- |
| Phobius | 0 | 0.5 | 1 | 0.850 | 0.740 | 0.516 | 0.842 | 0.578 |
| Philius | 0 | 0.5 | 1 | 0.842 | 0.741 | 0.516 | 0.800 | 0.553 |
| Topcons2 | 0 | 0.0 | 0.0 | 0.742 | 0.331 | 0.0 | 0.0 | 0.0 |
| PureseqTM | **1** | **1** | **1** | **0.908** | **0.978** | **0.806** | **0.833** | **0.758** |

#### 5ldwY

This protein is the chain Y from the mammalian respiratory complex I, which has 141 residues and 4 transmembrane helices. The most similar protein to 5ldwY in the training dataset has TM-score 0.545, which indicates that there is no strong structural similarity. As shown in Figure 5, our method predicted the correct number of transmembrane helices, while Topcons2 predicted one and Phobius failed to predict this protein as a TMP. Although Philius successfully predicted the correct number of transmembrane helices, in terms of residue-level accuracy, our method is significantly better than Philius (Table 4). In particular, the Q2, SOV and Mcc of our method is 0.823, 0.928, and 0.642, respectively, which is 6%, 5%, and 10% better than that of Philius. From this case, we may also find out that Philius tends to predict longer transmembrane segments, which will cause a better recall but a lower precision. Another interesting phenomenon from this case study is that although using a similar model architecture, say HMM in Phobius and DBN in Philius, it seems that for most of the cases Philius outperforms Phobius in terms of protein-level accuracy. This hypothesis could also be inferred from the fact that with the inclusion of DBN features to our model, the protein-level accuracy increases significantly.

**Figure 5.**
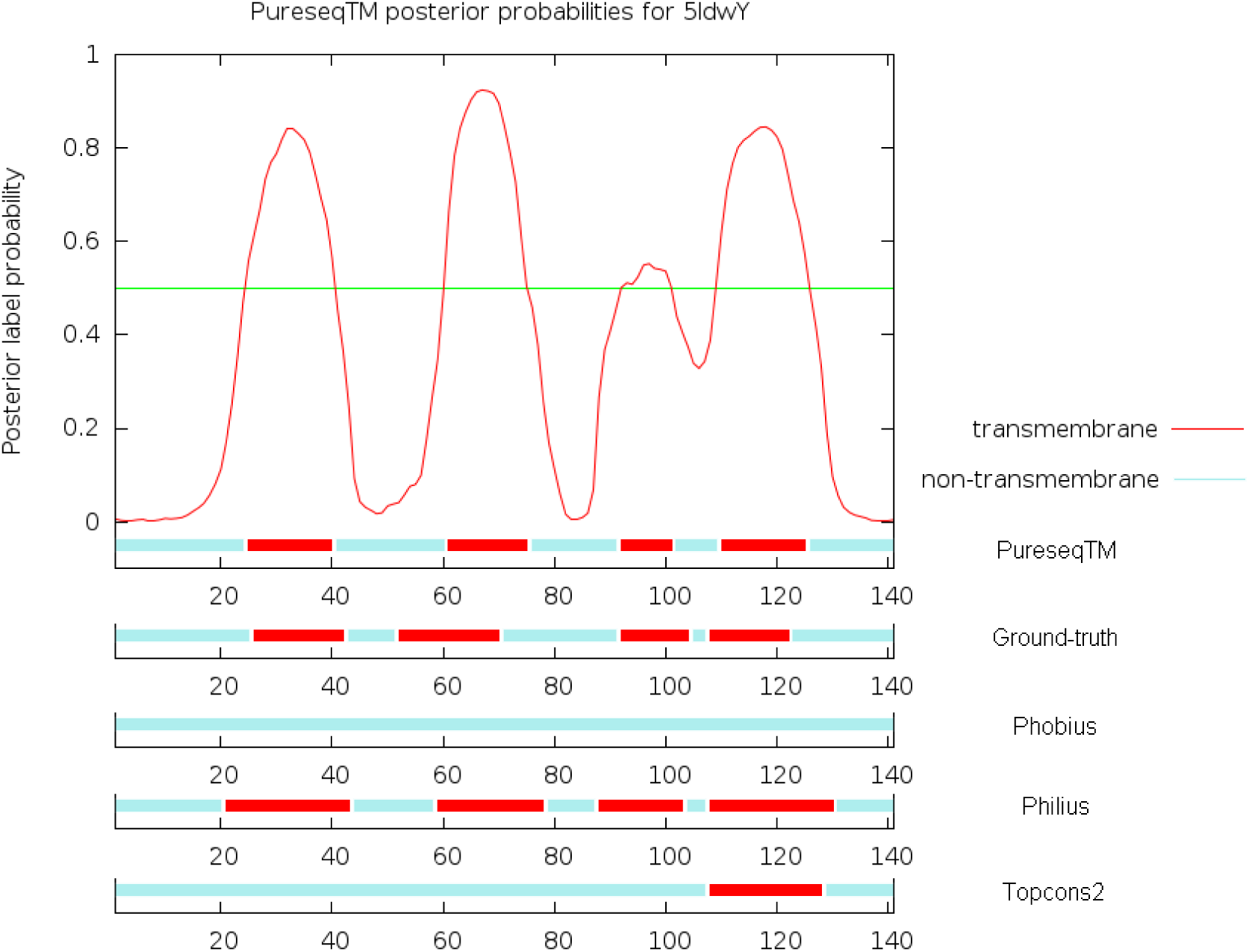
Case study of the transmembrane topology prediction of 5ldwY.

**Table 4.**
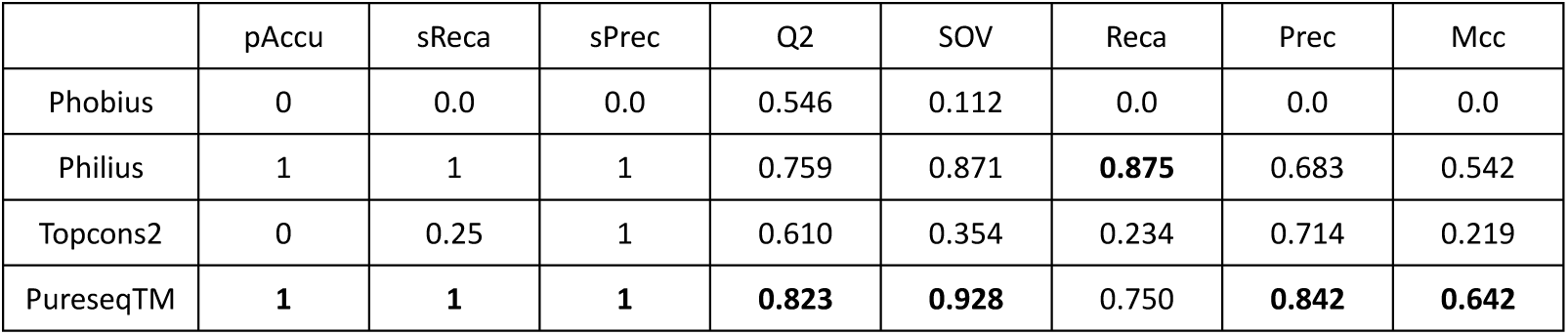
Transmembrane topology prediction accuracy of 5ldwY.

#### 4he8E

This protein is the chain E from the respiratory complex I from Thermus thermophiles, which has 95 residues and 3 transmembrane helices. This protein has structural analogs in the training dataset, where the most similar protein has TM-score 0.694 to this protein. The sequence homologs (measured by Meff) of this protein is 1004. Figure 6 shows that our method and Topcons2 predicted the correct number of transmembrane helices, while Phobius and Philius predicted two. In terms of residue-level accuracy, the profile-based consensus method Topcons2 outperforms PureseqTM, especially in SOV and Mcc (Table 5). This is normal that when a protein has a large amount of sequence homologs, the profile-based approach will gain increased performance in the residue-level measurements, as indicated in Table 1. Thus, if the profile model of PureseqTM is used for this case, we shall obtain a much better prediction than Topcons2 (Table 5). Specifically, the Q2, SOV and Mcc of our profile mode is 0.852, 0.983, and 0.708, respectively, which is 4%, 9%, and 3% better than that of Topcons2.

**Figure 6.**
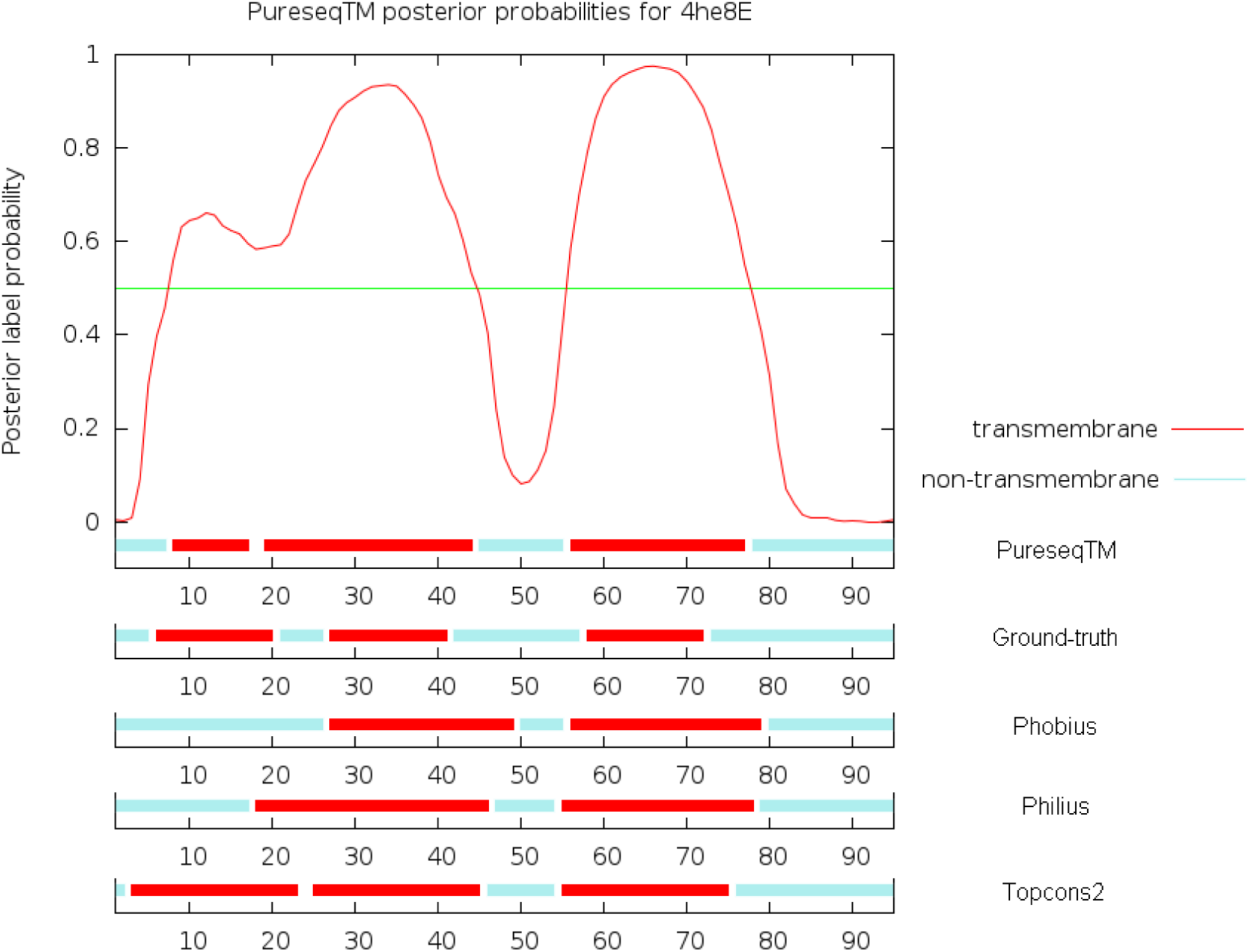
Case study of the transmembrane topology prediction of 4he8E.

**Table 5.** Transmembrane topology prediction accuracy of 4he8E.

|  | pAccu | sReca | sPrec | Q2 | SOV | Reca | Prec | Mcc |
| --- | --- | --- | --- | --- | --- | --- | --- | --- |
| Phobius | 0 | 0.667 | 1 | 0.663 | 0.669 | 0.667 | 0.638 | 0.326 |
| Philius | 0 | 0.667 | 1 | 0.663 | 0.678 | 0.733 | 0.622 | 0.335 |
| Topcons2 | 1 | 1 | 1 | 0.811 | 0.895 | **1.0** | 0.714 | 0.676 |
| PureseqTM | 1 | 1 | 1 | 0.800 | 0.802 | 0.933 | 0.724 | 0.628 |
| PureseqTM<sup>p</sup> | **1** | **1** | **1** | **0.853** | **0.968** | 0.933 | **0.792** | **0.717** |

### Discrimination of transmembrane and non-transmembrane proteins

Before predicting the transmembrane topology, a preliminary task is to distinguish TMPs and non-TMPs. This is critical for the application to genomic or proteomic data. To fulfill this task, we trained a model to discriminate TMPs and non-TMPs by randomly adding ∼1000 proteins from the non-TMP dataset to the training set. To test the performance of this discrimination model, we compared with Phobius, Philius and Topcons2 on the discrimination dataset that contains 440 TMPs (including both single-pass and multi-pass transmembrane proteins) and ∼6400 non-TMPs. We define true positives (TP) and true negatives (TN) as the numbers of correctly predicted TMPs and non-TMPs, respectively; whereas false positives (FP) and false negatives (FN) are the numbers of misclassified TMPs and non-TMPs, respectively.

As shown in Table 6, we observe that our method PureseqTM performs comparable to the other methods on this discrimination dataset, where all the four methods have relatively high success rate in distinguishing TMPs from non-TMPs. If we take a closer look at the 15 false negatives of PureseqTM, 12 (12) of them are also false negatives of Phobius (Philius), which indicates that the failure rate of discrimination of PureseqTM depends largely on the Phobius and Philius. Furthermore, among the 15 FNs, 8 of them (53.3%) are actually single-pass transmembrane proteins (sTMPs), which are not included in our training dataset. Similarly, if we take a look at the 50 false positives, 45 (31) of them are also false positives of Phobius (Philius). Among the 50 FPs, 35 of them (70%) are predicted to be sTMPs. These phenomena indicate that our discrimination model can be reliably applied to distinguish mTMPs from non-TMPs. However, there is still room for improving the success rate to discriminate sTMPs from non-TMPs.

**Table 6.**
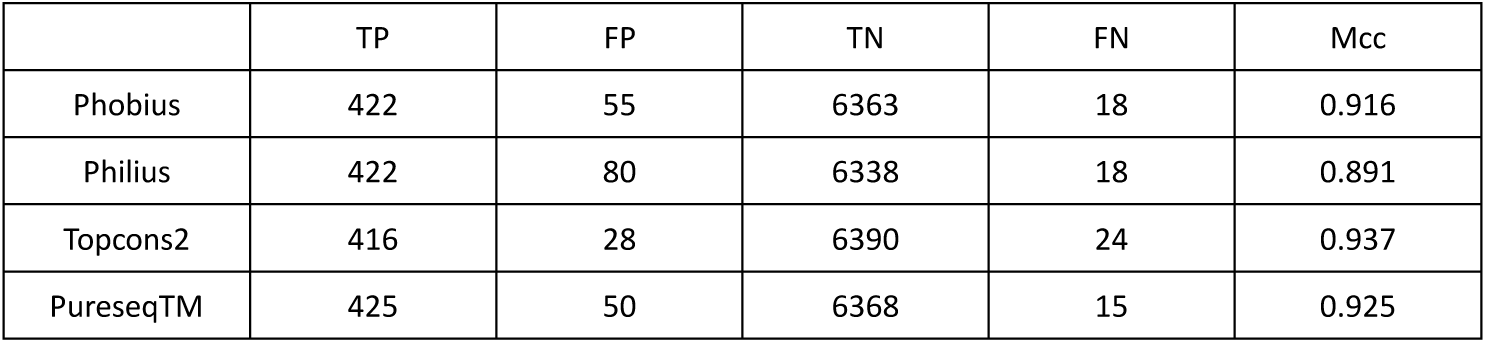
Discrimination accuracy of transmembrane and non-transmembrane proteins on the discrimination dataset.

### Application to Human proteome

Given the lower rate of false classifications of TMPs and non-TMPs, as well as the higher rate of correct identification of transmembrane segment, it should be interesting to see the application of PureseqTM for detecting the Human membrane proteome. To do so, we obtained the 20416 Human protein sequences from UniProt [24] to extract the reviewed transmembrane (TM) regions as ground truth, which results in total 5238 TMPs. Among these 5238 TMPs, 2399 are annotated as sTMPs and the remaining 2839 are mTMPs. It should be noted that even when there is proof for the existence of TM segments, it is difficult to determine their boundaries. Thus, only 52 of the TMPs has experimental evidence (i.e., UniProt ECO: 0000269), while most of the ground truth annotations are based on predictions by TMHMM, Memsat and Phobius. To show the performance of PureseqTM, we first ran a discrimination study on Human TMPs and non-TMPs, we then predicted the TM topology segments on each identified TMP. We compared our method with Phobius, Philius, and Topcons2, respectively. Finally, several case studies are shown to illustrate the usefulness of PureseqTM for (i) correcting the UniProt entries whose TM segments are mislabeled, and (ii) discovering novel mTMPs that are not annotated by UniProt.

Table 7 shows that PureseqTM correctly identifies 4912 (93.7%) TM proteins, second best only to Phobius that is one of the references for the ground-truth. Taking a closer look at the 326 false negatives of PureseqTM, we find that 251 (286) of them are also false negatives of Phobius (Philius). Interestingly, among the 326 FN, only 36 (11%) of them are mTMPs, whereas most of them belong to sTMPs. In particular, among the 290 sTMPs, about 115 (155) of them belong to type I (type II) signal-anchor sTMP, and about 20 of them belong to mitochondria TM proteins (mtTMPs). Therefore, more efforts shall be paid to improve the recognition rate for sTMPs and mtTMPs.

**Table 7.** Discrimination accuracy of transmembrane and non-transmembrane proteins on the Human proteome from UniProt.

|  | TP | FP | TN | FN | Mcc |
| --- | --- | --- | --- | --- | --- |
| Phobius | 4939 | 514 | 14879 | 299 | 0.898 |
| Philius | 4768 | 414 | 14708 | 470 | 0.886 |
| Topcons2 | 4812 | 466 | 14752 | 426 | 0.886 |
| PureseqTM | 4912 | 380 | 14852 | 326 | 0.910 |

Table 8 shows that PureseqTM reaches the second best to Phobius in terms of protein-level accuracy, segment-level accuracy as well as residue-level accuracy. This result implies that the TM topology boundaries detected by PureseqTM are closer to the UniProt annotation than those detected by Philius and Topcons2. However, it should be noted that the ground-truth itself in this task is actually a consensus annotation result, which will bias heavily to Phobius.

**Table 8.** Overall transmembrane topology prediction accuracy on the 5238 reviewed Human membrane proteins from UniProt.

|  | pAccu | sReca | sPrec | Q2 | SOV | Reca | Prec | Mcc |
| --- | --- | --- | --- | --- | --- | --- | --- | --- |
| Phobius | 0.734 | 0.916 | 0.915 | 0.936 | 0.935 | 0.857 | 0.810 | 0.785 |
| Philius | 0.651 | 0.885 | 0.867 | 0.934 | 0.922 | 0.830 | 0.772 | 0.750 |
| Topcons2 | 0.669 | 0.894 | 0.876 | 0.933 | 0.916 | 0.813 | 0.799 | 0.759 |
| PureseqTM | 0.696 | 0.920 | 0.888 | 0.934 | 0.932 | 0.803 | 0.818 | 0.759 |

Here we point out that the TM topologies of a variety of UniProt entries are not correctly annotated. Specifically, there exist 186 UniProt entries in which the number and boundaries of TM segments only match with Phobius, but not with Philius, Topcons, or PureseqTM (Supplemental Table S4). For example, sodium-dependent phosphate transport protein 3 (UniProt ID: O00624, and gene ID: SLC17A2) has 439 residues and was reported to have 6-12 TM segments [31]. Phobius and UniProt annotate 9 segments, but Philius, Topcons, and PureseqTM all predict 11 segments (Figure 7). An alternative evidence to show SLC17A2 being an 11 segment mTMP comes from the PredMP service [32] (http://database.predmp.com/#/databasedetail/O00624) that performs *de novo* folding assisted by the predicted contact map from RaptorX-Contact [33]. As shown in Figure 7, the Meff (number of non-redundant sequence homologs) value of SLC17A2 reaches ∼9,680, and ln(Neff) is 4.9 where Neff is the length-normalized Meff. According to literature [34], when ln(Neff) is larger than 3.5, the predicted 3D models by RaptorX-Contact on average will have TMscore⩾0.6, which indicates that this 3D model is likely to have a correct fold. Hence, we strongly believe that SLC17A2 is an mTMP with 11 TM segments, and the UniProt annotation is incorrect.

**Figure 7.**
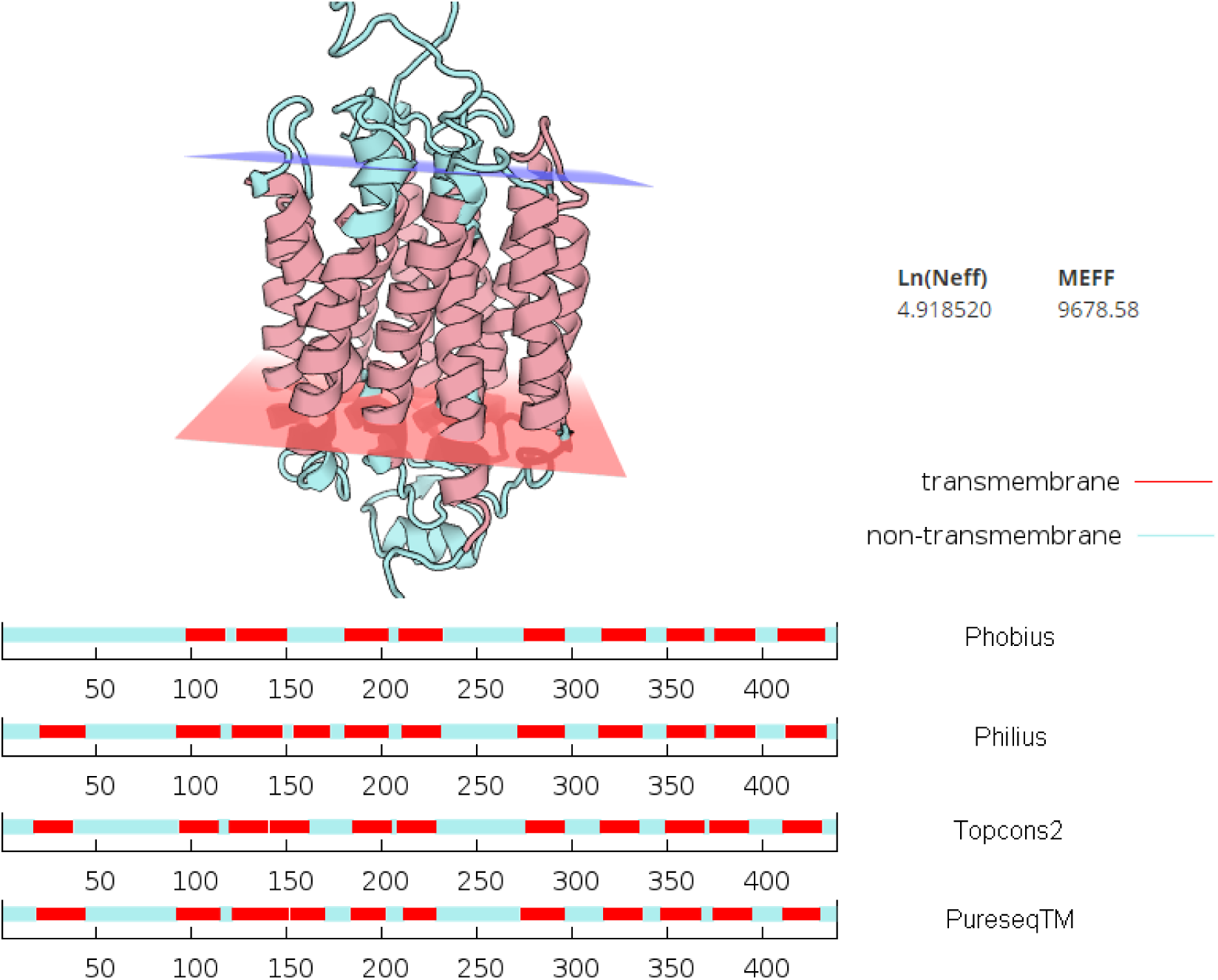
Case study of the transmembrane topology prediction of SLC17A2 (UniProt ID: O00624). The 3D structure model is *de novo* folded by the PredMP server that utilized the predicted contact map from RaptorX-Contact. The number of non-redundant sequence homologs (Meff) and log of length-normalized Meff (Neff) of this protein is ∼9,680 and 4.9, respectively.

Among the 380 false positives of PureseqTM, we found that 349 (209) of them are also false positives of Phobius (Philius) and 262 (68.9%) of them are predicted to be sTMPs. For the remaining 118 UniProt entries that are predicted to be mTMPs by PureseqTM, we figured out that 41 of them are also predicted to be a TMP by Phobius, Philius, and Topcons2 (Supplemental Table S5). This phenomenon indicates that these non-TMP UniProt entries might have a chance to be relevant to membrane (e.g., either buried within, interfacial to, or cross the membrane). We show here that at least the following 9 entries: Q9UMS5, Q8N3S3, Q9Y5W8, Q5JWR5, Q6NT55, P11511, P51690, Q9BXQ6, and O95197, satisfy these conditions according to the literature. For example, the putative homeodomain transcription factor 1 (UniProt ID: Q9UMS5, and gene ID: PHTF1) has 762 residues. In 2003, J. Oyhenart *et. al.* determined that PHTF1 should be an integral membrane protein localized in an Endoplasmic Reticulum (ER) [35]. Later on, Reta Birhanu Kitat *et. al.* performed an experiment based on high-PH reverse-phase StageTip fractionation to confirm that PHTF1 is a membrane protein [36]. According to our analysis, PHTF1 contains 7 transmembrane helices (Figure 8). This evidence is not only supported by the prediction result from Topcons2, but also from the published literature by A. Manuel *et. al.*, in which they proposed that the stretches of hydrophobic amino acid residues (from amino acids 99 to 115, 122 to 138, 477 to 493, 533 to 549, 610 to 626, 647 to 663, and 736 to 752) might be membrane-spanning domains [37]. The range of these transmembrane segments is quite close to PureseqTM and Topcons2 (Figure 8).

**Figure 8.**
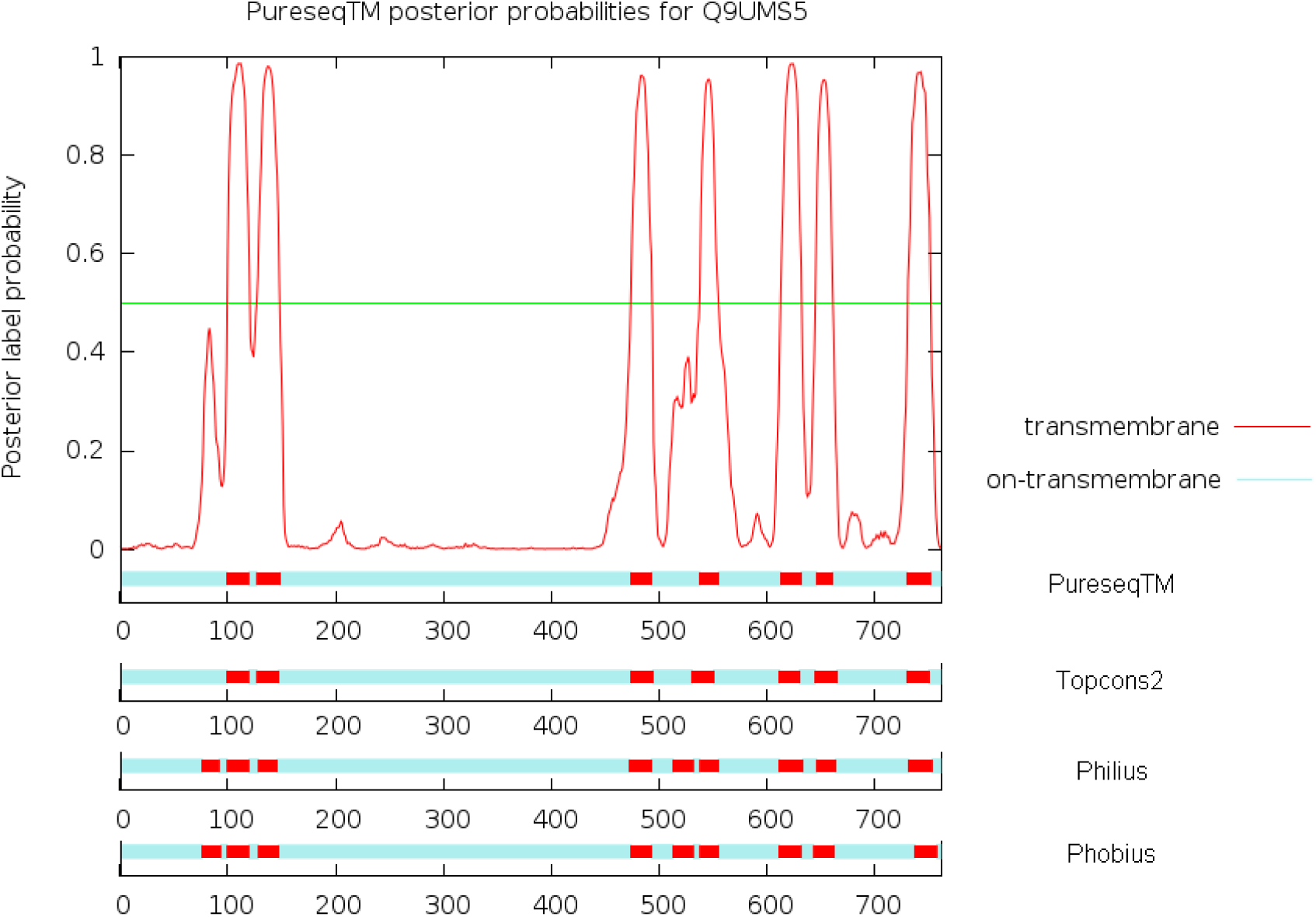
Case study of the transmembrane topology prediction of PHTF1 (UniProt ID: Q9UMS5).

Finally, we ask a question of whether or not PureseqTM could find a new membrane protein (MP) that neither is labeled by UniProt, nor is predicted to be a MP by Phobius, Philius, and Topcons2. To answer this question, we first collected a subset from the 118 UniProt entreis that are predicted to be non-TMP by all the other three methods. This list contains 11 entries, and at least one entry Q9BY12 (S phase cyclin A-associated protein in the endoplasmic reticulum, SCAPER) has strong literature evidence to be a membrane-related protein. Specifically, Tsang *et. al.* first reported that SCAPER is a perinuclear protein localized to the nucleus and primarily to the ER, in which SCAPER is most enriched in the membrane fraction [38]. Later on, Tsang *et. al.* further indicated that SCAPER may either be associated with the surface of the membrane or a transmembrane protein that spans the membrane at least twice [39]. This coincides with our prediction as shown in Figure 9.

**Figure 9.**
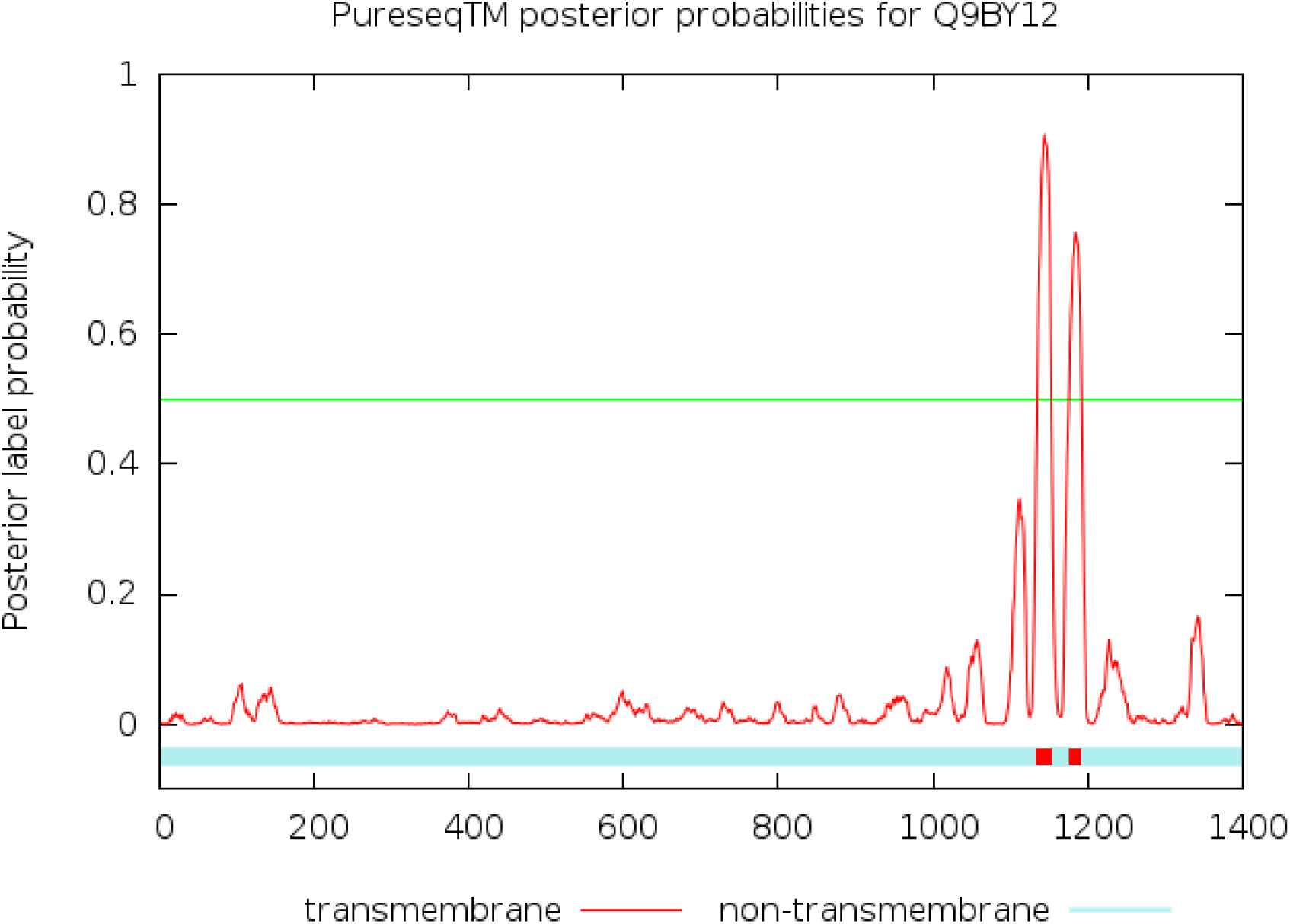
Case study of the transmembrane topology prediction of SCAPER (UniProt ID: Q9BY12).

### Runtime analysis

We test the execution time of PureseqTM, Phobius and Philius on a Fedora25 system with 128Gb memory and two E5-2667v4 (3.2 GHz) processors. As PureseqTM requires only amino acid sequence information, the runtime for a single protein with 500 residues is about 2.5 second, which is much faster than those profile methods that need at least several minutes (or even hours) to construct the MSA. When the protein length grows, the runtime increases linearly (Figure 10). By default, PureseqTM will first call Phobius and Philius to generate the HMM and DBN features. Therefore, it’s normal that the runtime of PureseqTM is roughly the summation of the runtime of Phobius and Philius. As shown in Table 1, the performance of PureseqTM does not differ much if the HMM and DBN features are not added. We denote this mode as PureseqTM_fast, which can accelerate the runtime considerably to less than 0.5 second for a 500 length protein (Figure 10).

**Figure 10.**
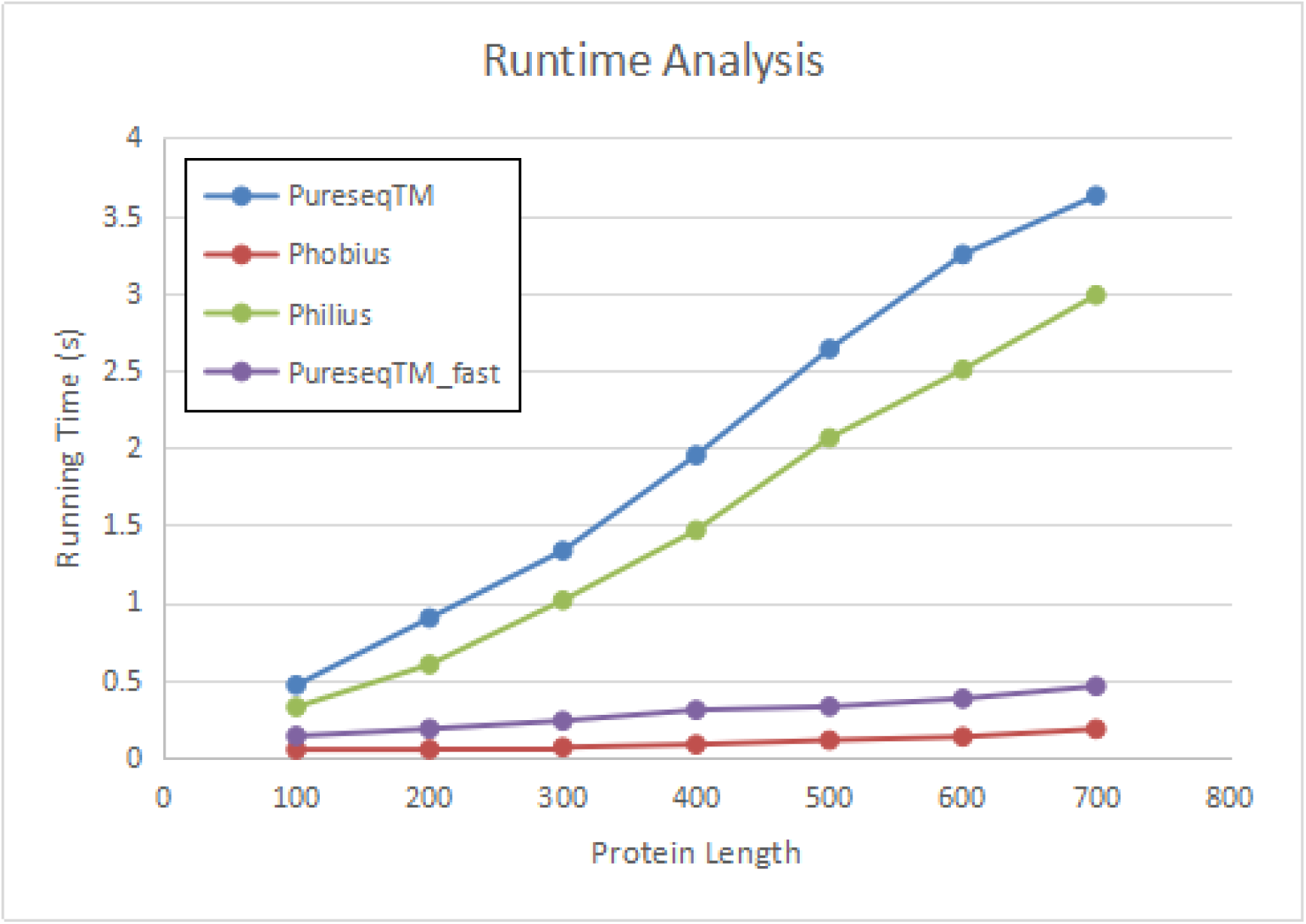
The runtime analysis of PureseqTM, Phobius and Philius with different protein lengths. Note that PureseqTM_fast is the fast mode of PureseqTM without using HMM and DBN features.

## Conclusion and discussion

In this paper, we proposed a deep learning method, PureseqTM, for transmembrane topology prediction from amino acid sequence only (i.e., pureseq features). PureseqTM makes its uniqueness from the other methods in that it employs a deep probabilistic graphical model, DeepCNF, to simultaneously capture long-range dependencies embedded in the input features as well as explicitly depict the interdependency between adjacent topology labels. Compared to HMM and DBN, CRF (a key module in DeepCNF) does not have the strict independence assumptions. This property allows CRF to accommodate any contextual information, which makes the feature design in CRF much more flexible than that in HMM and DBN. Experimental results show that PureseqTM performs much better than the state-of-the-art pureseq methods, and even reaches or outperforms the profile and consensus methods, in terms of protein-level, segment-level, and residue-level accuracy. Therefore, PureseqTM is the first approach, to our knowledge, that can reach high prediction accuracy while keeping efficient running speed, which enables the accurate annotation of the entire membrane proteome in reasonable time.

Our feature incremental study has three important findings. Firstly, the influence of the profile features in transmembrane topology (TM) is less important compared to that in other protein structural properties such as 3-state secondary structure element (SSE), 3-state solvent accessibility (ACC), and 2-state order/disorder region (DISO). According to the literature [14, 18], the difference of the prediction accuracy in terms of QX (here X indicates the number of labels, such as Q3 for SSE and ACC, and Q2 for DISO and TM) between pureseq features and profile features are 10%, 8.5%, 4%, and 2% for SSE, ACC, DISO, and TM, respectively. Secondly, we found that the HMM and DBN features could further improve the prediction accuracy with the pureseq features, but fail to do so with the profile features. This might explain the phenomenon that in some cases the prediction accuracy of a consensus approach does not outperform the single method integrated in that consensus approach. Thirdly, although PureseqTM incorporates the HMM and DBN features generated by the two state-of-the-art pureseq methods Phobius and Philius, we show that without these features (say, only use the one-hot amino acid encoding feature) PureseqTM can still reach higher performance than Phobius and Philius in terms of segment-level and residue-level accuracy.

One obvious limitation of our method is the relatively low performance when the input sequence is an sTMP, as shown in Table 6 and 7. This is normal because we excluded all the sTMPs from our training/validation data. The underlying reasons are two folds: (i) the physical-chemical properties of sTMPs are quite different from that of mTMPs, where the formers are mostly represented by water-soluble domains. If we add those sTMPs to train our model, the prediction accuracy of mTMPs will be influenced (data not shown); (ii) currently, compared to mTMPs, sTMPs only constitutes about 25% of the PDBTM database. However, according to our analysis, among ∼5200 human membrane proteins, ∼45% of them are sTMPs. This data imbalance issue will cause the trained model to be biased to mTMPs. Therefore, in order to solve this problem, we shall distinguish them separately by a discrimination model (as what we did for discriminating TMPs and non-TMPs), and train different models for predicting the TM region(s). Further, to solve the limited number of sTMPs in the PDBTM (less than 150 if we set the non-redundancy threshold to 25% sequence identity) for training purpose, there appears a Membranome database that provides structural and functional information about more than 6000 sTMPs from a variety of model organisms [40].

We may further improve the TM topology prediction for mTMPs by leveraging the 2D pairwise features embedded in the co-evolutionary information [33]. It is obvious that the pairwise TM helical-helical interactions are critical for the formation of TM topology, in which the underlying pairwise features are the 2D distance map that contains all square distances between each pair of the residues. Recently, these pairwise features, as described by contact map or distance map, could be predicted accurately from co-evolutionary analysis through ultra-deep learning models [33, 34, 41]. Recently, a similar approach has been successfully applied to predict the SSE and significantly improved the accuracy [42]. Thus, we believe that such approach could also improve the TM topology prediction by a large margin.

## Funding

This work was supported by the King Abdullah University of Science and Technology (KAUST) Office of Sponsored Research (OSR) under Awards No. FCC/1/1976-17-01, FCC/1/1976-18-01, FCC/1/1976-23-01, FCC/1/1976-25-01, FCC/1/1976-26-01, and URF/1/3450-01 to X.G. This work was also supported by National Institutes of Health (NIH) [R01GM089753] and National Science Foundation (NSF) [DBI-1564955] to J.X.

### Conflict of Interest

none declared.

## Method

### Datasets

There are three datasets used in this work: training, testing, and discrimination. We train and validate our method on the training dataset. The test dataset is applied for testing our method and comparing it with other approaches. The discrimination dataset is used for distinguishing transmbrane and non-transmembrane proteins.

#### Training dataset

The dataset used to train our proposed method is a subset of 510 transmembrane proteins created in Jul 2016 from PDBTM [11], in which any two proteins share less than 25% sequence identity [32]. Specifically, we choose 328 alpha-helical multi-pass transmembrane proteins (mTMPs) from this 510 dataset, and divide them randomly into two equally sized sets: one for training and the other for validation (Supplemental Table S1 and S2).

#### Testing dataset

To test the performance of our method and compare with other approaches, we collect a set of TMPs from PDBTM which are released after Jul 2016, or with no homology with the 510 dataset. To remove redundancy with the 328 training/validation dataset, we use a strict rule that (i) any two proteins in the test set share less than 25% sequence identity; (ii) there is no protein in the test set that shares >25% sequence identity or BLAST [43] E-value <0.001 with any proteins in the training/validation dataset. This creates a test dataset containing 39 mTMPs (Supplemental Table S3). We use a protein structure alignment tool DeepAlign [29] to perform 3D structure comparison between the proteins in the test dataset and those in the training dataset.

#### Discrimination dataset

To show the performance of discriminating transmembrane proteins (TMPs) and non-transmembrane proteins (non-TMPs), we collect a subset containing 440 alpha-helical TMPs from the 510 dataset as the TMP dataset. For the non-TMP dataset, we first download the PDB25 dataset released in Sep 2016 [44], in which any two proteins share less than 25% sequence identity. We then exclude the proteins in PDB25 sharing >25% sequence identity or having a BLAST E-value <1 with any of the 510 dataset. This results in 6418 proteins as the non-TMP dataset.

To label each residue from a given TMP sequence, we used the following 9 labels extracted from PDBTM [11]: 1 (Side1), 2 (Side2), B (Beta-strand), H (alpha-helix), C (coil), I (membrane-inside), L (membrane-loop), F (interfacial helix), and U (unknown localizations). As in this work we focus on alpha-helical mTMPs, the 9 labels are reduced to binary classification in which label H is denoted as ‘1’ and all other labels are denoted as ‘0’. For non-TMPs, we label all residues as ‘0’.

### Input features

Our method in the ‘pureseq’ mode only relies on the residue-related features. If evolutionary profile information is provided, our method in the ‘profile’ mode could also take as input the evolution-related features.

If the ‘pureseq’ mode is chosen, our method only takes input the 36 residue-related features: (i) amino acid identity represented as a binary vector of 20 elements (or, one-hot encoding); (ii) reduced AAindex [23] which contains 5 highly interpretable numeric patterns of amino acid variability (see Table 2 in [26]). These features allow a richer representation of amino acids that reflect polarity, secondary structure, molecular volume, codon diversity, and electrostatic charge [16]; (iii) 4 predicted probabilities of the transmembrane topology labels from Phobius [12], which are cytoplasmic (label ‘`i`’), non-cytoplasmic (label ‘`o`’), transmembrane region (label ‘`m`’), and signal peptide (label ‘`s`’), respectively; (iv) 7 predicted transition probabilities between the transmembrane topology labels from Philius [8], which are ‘`ii`’, ‘`oo`’, ‘`mm`’, ‘`ss`’, ‘`mi`’, ‘`mo`’ and ‘`xm`’, respectively, where ‘`x`’ indicates ‘`i`’ or ‘`o`’.

If the ‘profile’ mode is chosen, besides the 20 one-hot encoding and 5 AAindex features, we use additional 40 evolution-related features. In particular, we use 20 position specific scoring matrix (PSSM) generated by PSI-BLAST [43] to encode the evolutionary information at each residue. We also use 20 hidden Markov model (HMM) profile generated by HHpred [45], which is complementary to PSSM to some degree. The reason why we do not use the 4+7 predicted probabilities of the topology labels in the profile mode is due to the fact that they do not improve the prediction accuracy (Table 1), especially the protein-level accuracy, on the validation dataset. The evolution-related features could be generated as a TGT file using the procedure https://github.com/realbigws/TGT_Package.

### DeepCNF training

The training procedure for the topology prediction using the DeepCNF model is based on the procedure used for training protein secondary structure element (SSE) [15]. In particular, we fix the model architecture with the following parameters: 5 layers, 100 neurons and 11 window length per position for each layer.

Although similar with the model to train SSE, here we use two cascaded procedures to train the predictor for 2-state transmembrane topology: (i) a model to distinguish TMPs and non-TMPs, and (ii) a model for accurately detecting transmembrane topology regions. We describe the detailed training procedures for each of them.

#### Model for detecting transmembrane topology regions

We first describe how to train the model on mTMPs. We train our model using the 164 entries from the training set and validate our model on the 164 entries from the validation set. As the model architecture is fixed, there is only one tunable hyper-parameter lambda in DeepCNF model, which is used for reducing over-fitting with a L2 norm. We try a variety of values ranging from 0,0.5,1,1.5,2,2.5,3.5,5,7.5,10,12.5,15,17.5, 20,22.5,25,27.5,30 and find out that lambda=20 produces the best performance on validation dataset (actually, there is no large difference when lambda ranges from 5 to 30). Although the trained model could detect topology regions accurately, it does not performwell on the discrimination task. The underlying reason is that we do not feed any non-TMP as the training data. We denote this trained model as ‘`puretm.model`’. A similar approach could be applied for training the model with evolution-related features, and we denote this model as ‘`proftm.model`’.

#### Model to distinguish TMPs and non-TMPs

To train this model, we use the same 164 entries from the training set and randomly choose 1000 proteins from the non-TMP dataset, while keeping the 164 entries from the validation set as is. Note that there is a highly imbalanced non-TMP/TMP ratio (around 15:1) in this task. To deal with the imbalanced distribution of the topology labels, we train the DeepCNF model by maximizing AUC which is an unbiased measure for class-imbalanced data [46]. An alternative approach to train this discrimination model is to initialize the DeepCNF with the trained model ‘`puretm.model`’, and to early stop when the AUC value on the validation set declines. We denote this trained model as ‘`detect.model`’.

#### Feature incremental study

To show the importance of each type of features in the ‘pureseq’ mode, we conduct an incremental study which is similar to a reverse operation of ablation study. Specifically, as there are four types of features: one-hot encoding, AAindex, HMM, and DBN, we incrementally add them to train the DeepCNF model on the 164 training set and check the performance on the 164 validation set. The order we choose for the four types of features is according to their model complexity. If the newly added feature type could significantly improve the prediction accuracy on the validation dataset in all measurements, then we conclude that such type of feature will make large contribution to the prediction. Otherwise, the feature might not be that important. A similar feature incremental study could be conducted for the ‘profile’ mode.

### Transmembrane topology prediction

When an amino acid sequence is given, PureseqTM will first call `detect.model` to distinguish TMP and non-TMP (Figure 1). If TMP is identified, then PureseqTM will employ `puretm.model` to predict the transmembrane topology (‘1’ for transmembrane and ‘0’ for non-transmembrane) and also the corresponding probabilities at each residue. Our method also enables the sequence profile as the input when the ‘profile’ mode is on. It should be noted that if the predicted transmembrane segment satisfies the two conditions: (i) the segment length is above 30 and (ii) there exist two peaks in the predicted probability, then PureseqTM will cut this segment into two at the position where the local minimum of the probability is found.

### Programs to compare

We compare our method with the following programs: Phobius [12], Philius [8], and Topcons2 [13] for 2-state transmembrane topology prediction. Phobius and Philius are methods that only rely on the input sequence information (i.e., ‘pureseq’ methods); Topcons2 is a consensus approach that combines the outputs from different predictors ranging from Phobius, Philius, SCAMPI [47] and OCTOPUS [10]. It should be noted that SCAMPI is a pureseq method, whereas OCTOPUS relies on the evolutionary profile (i.e., a ‘profile’ method).

We run Phobius and Philius with their default parameters. For Topcons2, we submit all the relevant sequences from the datasets to its server to obtain the prediction results. It should be noted that all of the three approaches return more labels (such as signal peptide) other than binary topology prediction. To transfer their results into 2-state transmembrane topology, we only keep the probability in state ‘transmembrane’ as label ‘1’, while regarding other states as label ‘0’ (i.e., non-transmembrane).

### Data and software availability

The web server implementing the DeepCNF model for transmembrane topology prediction from the amino acid sequence features only is publicly available at http://pureseqtm.predmp.com/. The code for the stand-alone package is maintained on GitHub (https://github.com/PureseqTM/pureseqTM_package). The list of the training, validation, and testing dataset is available in the Supplemental Material. For more detailed information about the relevant dataset, such as the Human proteome dataset, users are suggested to visit https://github.com/PureseqTM/PureseqTM_Dataset.

## Supplemental Material

### S1 Segment OVerlap (SOV) Score

The Segment Overlap score (SOV) measures overlap between the observed and the predicted transmembrane (TM) topology segments instead of per-residue accuracy. The predictions that have high per-residue accuracy but deviate from experimental segment length distributions have lower SOV scores. SOV score ranges from 0 to 1 with 1 indicating the perfect overlap.

Brief description of SOV is as follows. To calculate 2-state SOV, the predicted transmembrane topology of one protein sequence is parsed into segments such that each segment has a single topology type (either TM:1 or non-TM:0). Let S1 be the observed transmembrane topology and S2 the predicted transmembrane topology. For each type, *i* ∈ {0,1}, *S*(*i*) is the set of segment pair (*s*1, *s*2) with type *i* where *s*1 is from S1, *s*2 is from S2, and there must be at least one residue overlap with *s*1 and *s*2. That is, *S*(*i*) = {(*s*1, *s*2): *s*1 ∩ *s*2 ≠ 0, *s*1 *and s*2 have type *i*}. In contrast, *S*′ (*i*) = {*s*1: *s*1 ∩ *s*2 = 0, *s*1 *and s*2 have type *i*}.

Then the segment overlap score between S1 and S2 is calculated as follows:

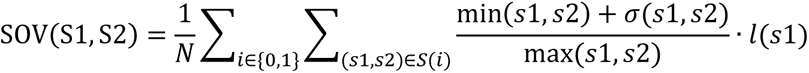

where min(s1, s2) is the length of the overlap between s1 and s2, max(s1, s2) is the length of the total span of s1 and s2, and *l*(s1) is the length of s1, *σ*(s1, s2) is defined as min(max(s1, s2) − min(s1, s2), min(s1, s2), 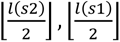) and *N* is defined as 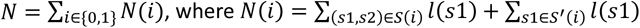

### S2 Dataset of training/validation and testing

**Supplemental Table S1.**
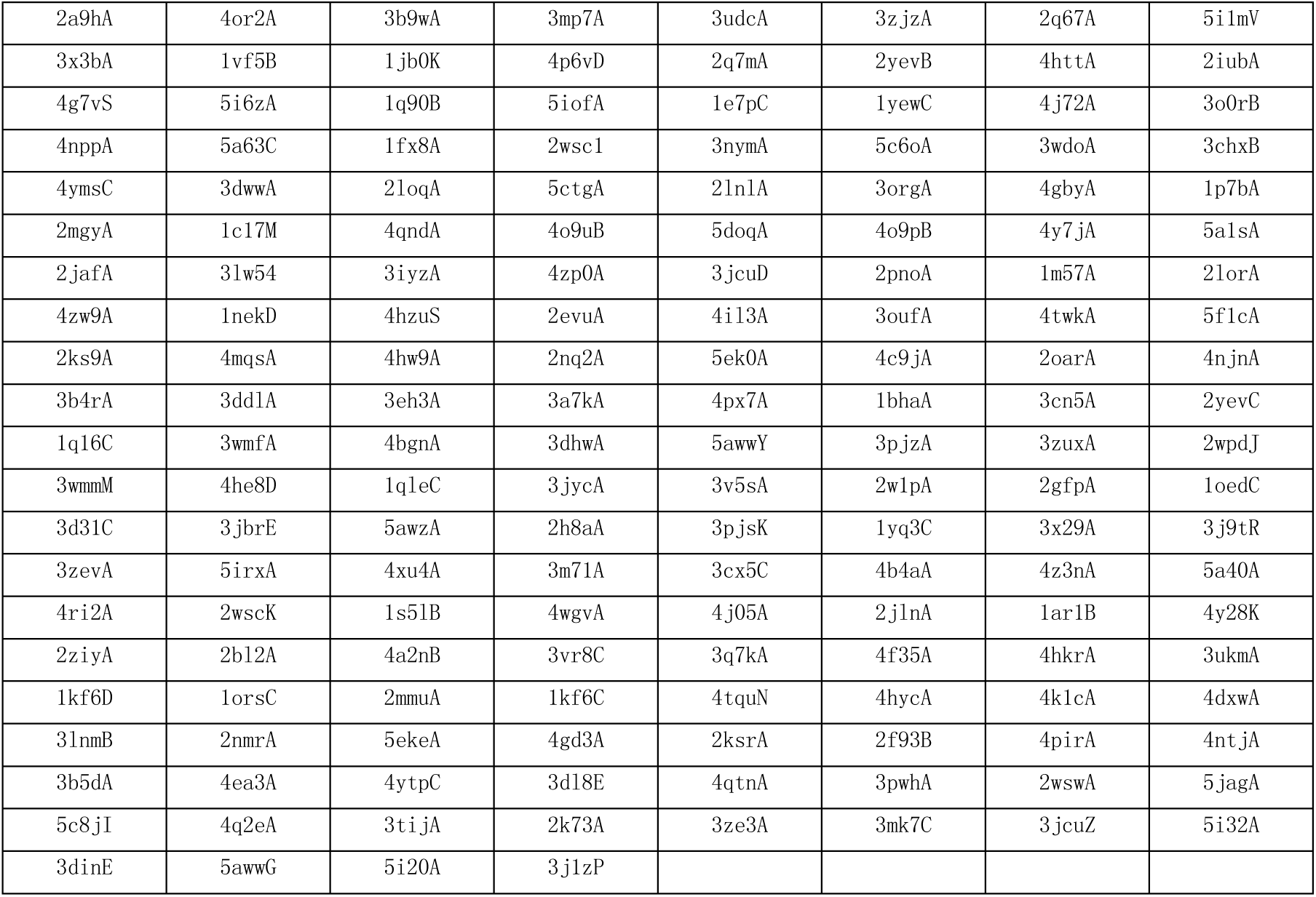
A list of 164 training data.

**Supplemental Table S2.** A list of 164 validation data.

|  |  |  |  |  |  |  |  |
| --- | --- | --- | --- | --- | --- | --- | --- |
| 4atvA | 2akhA | 1fftB | 4xxjA | 3kj6A | 3effK | 2n4xA | 4knfA |
| 4rfsS | 3wo7A | 2yiuA | 2d57A | 4us3A | 3chxC | 4tq3A | 4kt0K |
| 3o7pA | 3um7A | 4o6yA | 4pgrA | 2kseA | 2z73A | 2xq2A | 3111A |
| 4gycB | 1fftC | 4gx5A | 4bwzA | 5iwsA | 4u4tA | 3jcuS | 3k3fA |
| 4o6mA | 4cSkA | 1yq3D | 4h33A | 4he8C | 5aymA | 4uc1A | 4y28G |
| 3j08A | 2lckA | 4bpmA | 3qe7A | 1kqfC | 4hyoA | 2f95B | 1x14A |
| 4ryiA | 2k9pA | 5abbZ | 4in5L | 1p49A | 5azbA | 3ejzA | 4nykA |
| 4huqT | 4r1iA | 5a6eB | 4ymkA | 1pw4A | 4zr0A | 2kyhA | 5i6cA |
| 4mbsA | 2m67A | 1gzmA | 2135A | 3iz1A | 4cadC | 4chvA | 4dojA |
| 2y5yA | 2ksdA | 4kppA | 1h2sB | 4bemJ | 5d0yA | 5ezmA | 4ltoA |
| 1nekC | 3vr8D | 2m6bA | 4od4A | 5araW | 3tx3A | 4rngA | 4cfgA |
| 5gar0 | 2wsc3 | 2bg9A | 2nr9A | 2mpnA | 2akhB | 5id3A | 2wscG |
| 3wxvA | 4tquM | 3vouA | 4ezcA | 4jkvA | 4dveA | 4m58A | 3gi8C |
| 3kp9A | 4o9pA | 2lomA | 3zk1A | 5fn2B | 4czbA | 2ksfA | 3rkoA |
| 4djiA | 3qnqA | 3p5nA | 4ytpD | 4z7fA | 4huqS | 3jcuR | 4qncA |
| 4l6rA | 4tkrA | 4zr1A | 3wvfA | 4p6vE | 5cfbA | 4p79A | 2r6gG |
| 3hd6A | 3mktA | 4rdqA | 4phzA | 2vpwC | 4mndA | 3mk7A | 5dirA |
| 3ux4A | 5araT | 2lp1A | 4rp8A | 4gluA | 4kjrA | 4xnvA | 4z34A |
| 2losA | 4quvA | 4p6vB | 5doqB | 3s0xA | 2r6gF | 4y28L | 3uq7A |
| 3ug9A | 5ee7A | 4f41A | 4wd7A | 4ky0A | 2llyA | 1xioA | 2wscL |
| 3v2wA | 4u91A | 2xutA | 4v1fA |  |  |  |  |

**Supplemental Table S3.** A list of 39 testing data.

|  |  |  |  |
| --- | --- | --- | --- |
| 1pb4D | 4m64B | 5ldwN | 5tcxA |
| 1zc7A | 5eikA | 5ldwY | 5tsaA |
| 2gfzA | 5fgnA | 5lnko | 5ttaA |
| 2m0qA | 5h1qA | 5mg3F | 5voxc |
| 2mn6B | 5ijiA | 5mrwA | 5vreA |
| 3g6bA | 5l75F | 5n6hA | 5wtrA |
| 3rkoK | 5l75G | 5o0tA | 5wufA |
| 3rkoN | 5ldwJ | 5sv0B | 5x5yF |
| 4he8E | 5ldwK | 5sv9A | 5x5yG |
| 4j7cK | 5ldwM | 5t77A |  |

### S3 Dataset relevant to UniProt Human proteome

**Supplemental Table S4.** A list of 186 UniProt entries in which the number and boundaries of transmembrane segments only match with Phobius, but not Philius, Topcons, and PureseqTM.

|  |  |  |  |  |  |  |  |  |
| --- | --- | --- | --- | --- | --- | --- | --- | --- |
| Q9HDB5 | Q99572 | Q8IZS8 | Q8IV31 | Q9NVA4 | Q9HCM3 | Q8NGS6 | Q8NGL7 | 075352 |
| Q9HBU9 | Q6ZT12 | Q86VU5 | Q9NRX5 | Q6NXT6 | Q9Y6K0 | Q8NGS4 | Q9UGF6 | Q9NZP5 |
| 015120 | Q5T4S7 | Q5TH69 | Q16842 | Q16635 | Q5JZY3 | Q8NGT2 | Q8NGG0 | Q8NGE3 |
| Q8TEB9 | Q9P2D8 | Q9UIR0 | Q11203 | Q8IUA7 | Q15884 | Q68CR7 | Q8NG80 | Q8NGX3 |
| Q53FV1 | Q16572 | Q96MX0 | Q9BYT1 | Q96SE0 | Q9N2K0 | P60507 | P0C7N8 | Q8NGY0 |
| Q9BZA8 | A0A1B0GTY4 | P0C7U3 | Q96G79 | Q9UKB5 | Q9H1C3 | Q5T6L9 | B2RTY4 | Q8NGS5 |
| 075452 | Q6ZS81 | Q86YA3 | Q99624 | P08910 | P60509 | Q9NWS6 | 060431 | Q8NGT0 |
| Q13423 | Q9UBH6 | Q11130 | Q96JT2 | Q3MIX3 | Q76MJ5 | A6NC97 | Q8NH93 | P0C617 |
| 000624 | Q6ZU64 | 060774 | Q7Z3Q1 | Q9UM73 | B6SEH8 | Q5JX69 | Q8NGP3 | Q6IF42 |
| Q86XR5 | P98187 | Q96CU9 | Q8TBB6 | Q7Z5J8 | P05981 | A0A1B0GTK4 | A6NJW4 | Q8WUY8 |
| P58400 | A0A087X1C5 | 043464 | Q8NG04 | 095870 | Q5VW36 | Q8NGS9 | 014524 | Q8IW70 |
| Q9H209 | Q9NRD9 | Q9Y4D8 | P50443 | Q9NP78 | Q8NBF6 | Q8NG92 | Q86X29 | Q9P283 |
| C9JH25 | Q86SJ6 | Q8NCG7 | A0AV02 | Q4ZG55 | Q58HT5 | Q9H210 | Q9Y693 | Q14656 |
| Q9H208 | Q9Y2G3 | Q6ZUT9 | A6NIM6 | 043292 | Q96M19 | Q8WZ84 | Q96KK3 | Q5W0B7 |
| Q6J4K2 | Q8IWY9 | Q9UBM7 | Q9H1N7 | Q3MIP1 | Q07812 | 095047 | A6NDP7 | Q8N661 |
| Q7L5N7 | Q9P2K9 | Q9BS91 | Q96RN1 | Q6P9B9 | Q16611 | Q8NGM8 | Q8NH01 | Q96DC7 |
| Q86VR2 | Q8N6G5 | P61009 | Q96HH6 | P34903 | A6QL63 | Q6IEY1 | P30954 | Q8TDW4 |
| P42702 | Q7Z6A9 | P60602 | Q9BVT8 | 095461 | Q6UXD1 | Q8NGC5 | Q8NHC4 | Q96NR3 |
| Q86WI0 | P30988 | Q9H4F1 | Q9C0B7 | Q9NR82 | P03891 | P47890 | Q8NGS8 | Q9HCF6 |
| Q9H490 | Q8IU99 | 075204 | Q8N4L1 | Q96RP8 | Q8NGR1 | Q8NH00 | 095013 | 043173 |
| Q8NCS4 | Q96HV5 | Q7Z402 | Q15035 | Q6ZP80 | Q6UWJ1 |  |  |  |

**Supplemental Table S5.**
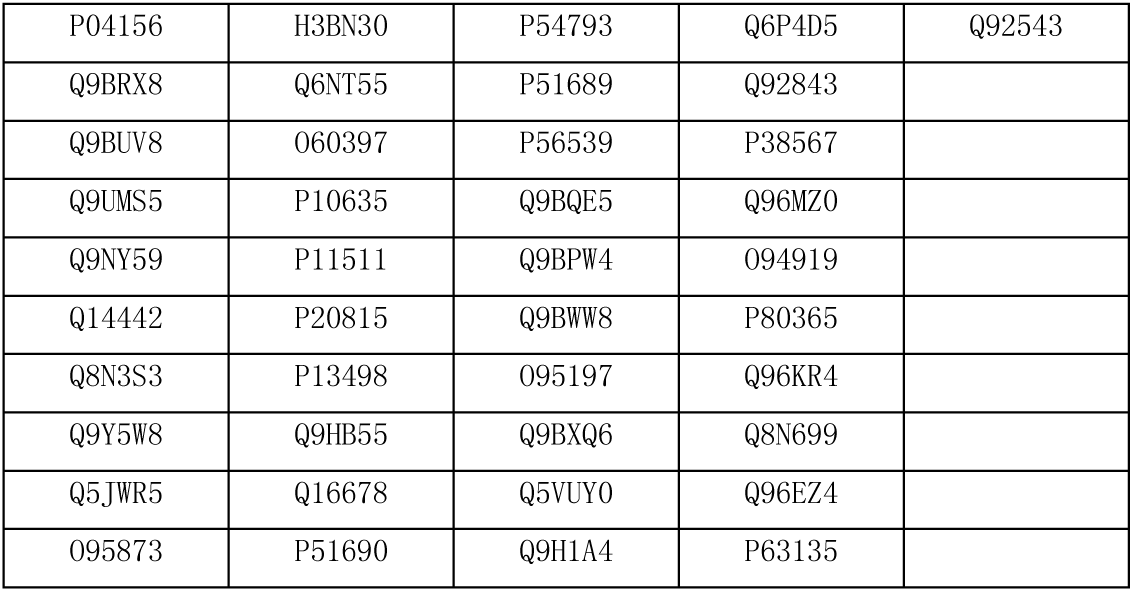
A list of 41 UniProt entries annotated as non-transmembrane protein but predicted to be a transmembrane protein by Phobius, Philius, Topcons2 and PureseqTM.

|  |  |  |  |  |
| --- | --- | --- | --- | --- |
| P04156 | H3BN30 | P54793 | Q6P4D5 | Q92543 |
| Q9BRX8 | Q6NT55 | P51689 | Q92843 |  |
| Q9BUV8 | 060397 | P56539 | P38567 |  |
| Q9UMS5 | P10635 | Q9BQE5 | Q96MZ0 |  |
| Q9NY59 | P11511 | Q9BPW4 | 094919 |  |
| Q14442 | P20815 | Q9BWW8 | P80365 |  |
| Q8N3S3 | P13498 | 095197 | Q96KR4 |  |
| Q9Y5W8 | Q9HB55 | Q9BXQ6 | Q8N699 |  |
| Q5JWR5 | Q16678 | Q5VUY0 | Q96EZ4 |  |
| 095873 | P51690 | Q9H1A4 | P63135 |  |

## Notes

#### Summary of Updates

Add Figure 10: The runtime analysis of PureseqTM, Phobius and Philius with different protein lengths.

https://github.com/PureseqTM/PureseqTM_Dataset

